# Atom-level mechanism of tapasin-independent peptide editing by Major Histocompatibility Complex class I molecules

**DOI:** 10.1101/2025.06.20.660712

**Authors:** Steven Turner, Rachel Darley, Jonathan W. Essex, Malcolm J. W. Sim, Andy van Hateren, Tim Elliott

## Abstract

Peptide binding to major histocompatibility complex class I molecules (MHC-I) and their presentation to cytotoxic immune cells is a keystone of the adaptive immune system. The selection of MHC-I bound peptides is facilitated by the chaperone tapasin, which allows MHC-I to iteratively sample peptides until they are loaded with optimal binding peptides, known as peptide editing. However, some MHC-I allotypes can select high affinity binding peptides independently of tapasin, and the molecular mechanism(s) for such peptide editing are unknown. Here, we used enhanced sampling molecular dynamics simulations of peptide-deficient MHC-I to investigate tapasin-independent peptide editing. Our simulations revealed transient disruption of hydrogen bonds between MHC-I and the peptide backbone could allow for peptide editing, a process we term "active displacement". Destabilisation of interactions with the peptide backbone, necessitates sequence-specific sidechain interactions to maintain peptide binding. Our active displacement model predicts surface expression levels for multiple MHC-I allotypes and accounts for the presentation of an immunogenic mutant KRAS-G12D neoepitope by HLA-C*08:02, but not by closely related HLA-C*05:01. Together our data provide a molecular mechanism for tapasin-independent MHC-I peptide editing, influencing the surface immunopeptidome and anti-tumour immunity.

**Significance Statement:** Major histocompatibility complex class I molecules (MHC-I) bind and present peptides to specialised killer cells of the immune system. These immune cells can unleash their cytotoxic effector functions if they recognise the peptide-MHC-I complex. Which peptides are presented by MHC-I is therefore highly important. Peptide selection is usually assisted by the tapasin protein, although some MHC-I molecules can select peptides independently of tapasin, but it is not known how this occurs. Here, we provide an atomistic description of the tapasin-independent peptide selection mechanism. Our mechanism applies to multiple MHC-I allotypes and illustrates how an immunogenic peptide is presented by one MHC-I molecule, but not by another closely related molecule. This new insight provides a rational basis for therapeutic treatments.

## Introduction

Major histocompatibility complex class I molecules (MHC-I) are cell surface receptors that bind peptides, primarily derived from endogenous protein catabolism, which they display at the cell surface to circulating CD8+ cytotoxic T-lymphocytes and natural killer (NK) cells. This process is critical for the immunosurveillance and detection of viral infection and cancer-associated mutations (1, 2). MHC-I is highly polymorphic (3), with MHC-I allotypes binding diverse repertoires of peptide ligands, typically 8-11 amino acids (4) and focused around the two or three “anchor” positions (5–8). This variability in MHC-I sequences and peptide repertoires makes prediction of peptide-MHC-I cell surface expression challenging (8–12).

MHC-I peptide loading occurs within the endoplasmic reticulum with the TAP1/2 heterodimer suppling a diverse pool of peptides (13–15). The relatively slow off-rates of low and medium affinity peptides (minutes-hours) is insufficient for MHC-I and β_2_-microglobulin (β_2_m) heterodimers to effectively sample peptides. Instead, MHC-I peptide sampling is orchestrated by tapasin, which recruits MHC-I-β_2_m to multi-subunit peptide-loading complexes (PLC) centred upon TAP1/2, with one or two editing modules comprising tapasin, ERp57, calreticulin and MHC-I-β_2_m (16–19). Within the PLC, tapasin facilitates iterative peptide exchange (peptide editing), skewing the repertoire towards high affinity peptides, enabling MHC-I to persist on the cell surface (20–23). Once a peptide is bound, peptide-loaded MHC-I heavy chain-β_2_m (pMHC-I) molecules dissociate from the PLC (16) and progress through the secretory pathway where they may engage in further editing by TAPBPR (24–26), before targeting to the plasma membrane (27), or recycling back to the ER in the case of insufficient peptide binding affinity (28–32).

While tapasin enhances peptide selection for all MHC-I allotypes, there is variation in the extent MHC-I allotypes engage tapasin (33–35). Some MHC-I allotypes, such as HLA-B*44:05, can effectively select high affinity peptides in the absence of tapasin, while tapasin is essential for HLA-B*44:02 (22, 35). The diversity in tapasin-dependence is likely to be evolutionarily selected (36), ensuring MHC-I antigen presentation in instances of tapasin/PLC downregulation, commonly observed through cancer-associated mutation (1, 37) or pathogen inhibition (38–40). Tapasin-independent peptide editing is less efficacious than tapasin-mediated editing (22, 41, 42), and results in a broader, more diverse peptide repertoire eliciting a greater number of immune responses (33, 43–45).

Here, we performed enhanced sampling molecular dynamics simulations on a panel of peptide deficient MHC-I allotypes with varying tapasin-dependency. Prior simulation strategies failed to generate predictive models of tapasin-independent peptide editing which generalise to multiple MHC-I allotypes. These models were restricted by limited sampling, focused on a few allotypes, or restricted dynamics to modelling of the peptide bound state (46–52). In this study, we first compared the dynamics of the HLA-B*44:02 and HLA-B*44:05 allotypes, which differ by a single polymorphic amino acid (HLA-B*44:02: Asp116; HLA-B*44:05: Tyr116) and in their dependence upon tapasin for cell surface expression (22, 35). We identified a dynamic signature that we termed active displacement, that we hypothesised could drive tapasin-independent peptide editing. Using our panel of MHC-I allotypes, we found all MHC-I allotypes that can independently self-edit peptide repertoire had an identical or mechanistically consistent dynamic signature, that was not found in their tapasin-dependent counterparts. We verified this hypothesis by mutating HLA-B*44:05 to disable the active displacement mechanism and found that the mutation impaired independent peptide editing ability. Last, we showed that active displacement accounts for the presentation of an immunogenic neoepitope by HLA-C*08:02, but not by closely related HLA-C*05:01, exemplifying how active displacement modulates immunogenic immune responses.

## Results

### The active displacement mechanism of independent peptide editing

To provide a mechanistic basis for independent peptide editing ability, we used molecular dynamics simulations with REST2 enhanced sampling to compare the HLA-B*44:02 and HLA-B*44:05 allotype pair in the peptide-deficient state. REST2 simulations enable the sampling of longer timescale dynamics that are inaccessible through conventional MD simulation. To visualise the simulation data, we plotted the distances between the α helices above the C and F pockets (figure 1a+b) and observed that both allotypes rapidly transitioned from an “open” crystal-structure like conformation (figure 1b+1c, upper) into a “closed” conformation (figure 1b+1c, centre), as expected to resolve interactions left in deficit by the missing peptide. In the closed conformation a hydrogen bond was often observed spanning the peptide binding groove between Asn77 and Trp147 (figure 1d, left). HLA-B*44:02 remained in this conformation for the remainder of the simulation time. By contrast, HLA-B*44:05 uniquely adopted a “hyper-closed” conformation (figure 1b+1c, lower) in which the C-pocket distance decreased from 16 Å to 14 Å (figure 1b), facilitated by the Trp147 sidechain rotating 29.3° with the indole nitrogen flipping towards the bottom of the peptide binding groove (figure 1d and supplementary figure 1). The simulations showed the reorientation of Trp147 was not a consequence of Trp147 sharing a hydrogen bond with either Tyr116 in HLA-B*44:05 or Asp116 in HLA-B*44:02, however, in 18.16% of simulation frames, we observed that Tyr116 in HLA-B*44:05 would transiently hydrogen bond to Asn77 (figure 1d, centre). The hydrogen bond between Tyr116 and Asn77 pulled Asn77 into a rotational orientation that was not conducive to hydrogen bonding to Trp147. The indirect destabilisation of Trp147 in HLA-B*44:05 allows the reorientation of its sidechain, and HLA-B*44:05 transitions into the hyper-closed conformation (figure 1c+1d, right). In comparison, an equivalent interaction between Asp116 and Asn77 was not observed in HLA-B*44:02, so Trp147 was not destabilised, and the hyper-closed conformation was not observed.

**Figure 1.**
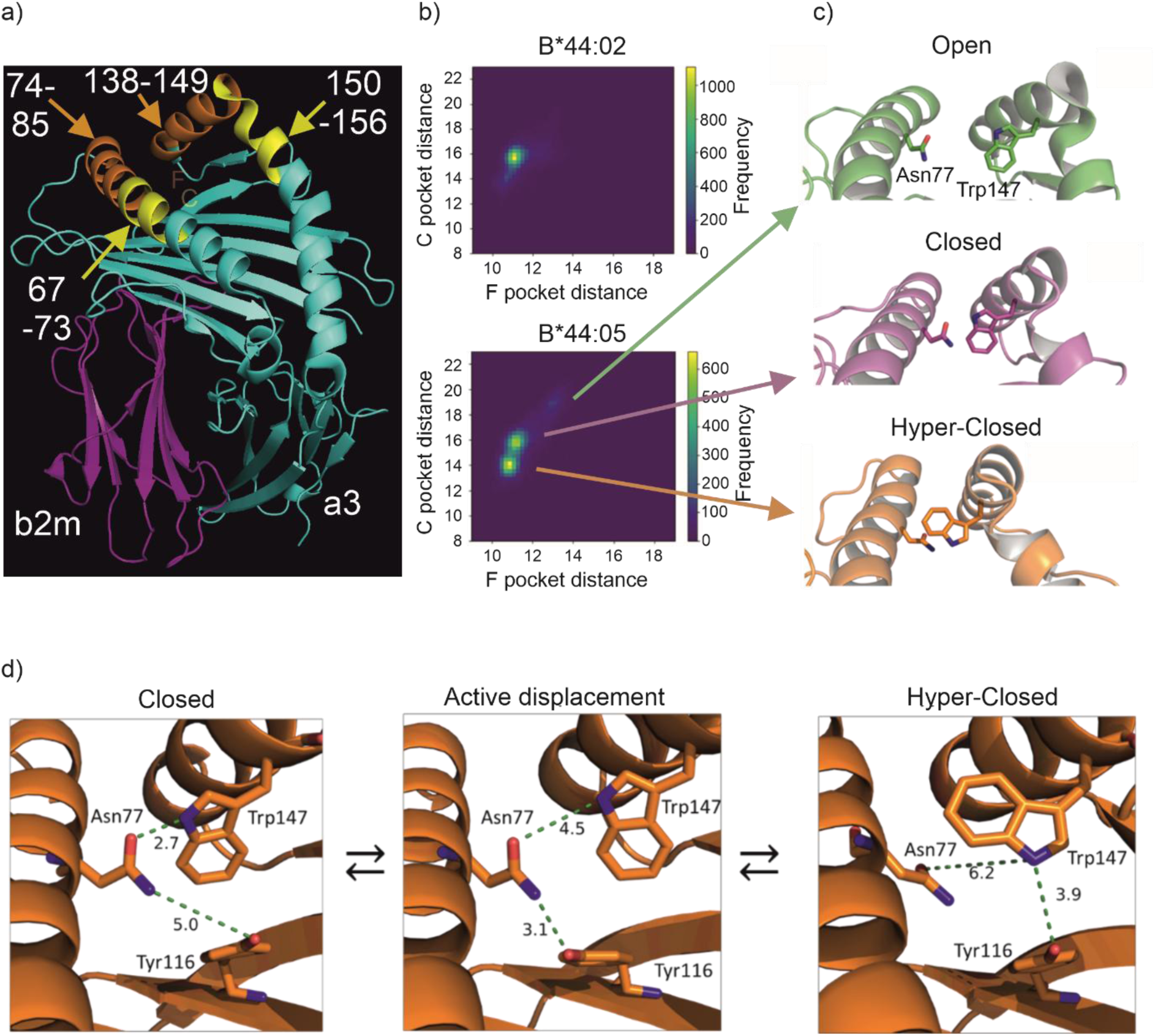
The HLA-B*44:05 active displacement mechanism. a) Cartoon of the HLA-B*44:02 structure (PDB: 3KPM) with structural annotations. The α2 helix can be subdivided into the α2-1 sub-helix (138–149), α2-1 sub-helix (150–159) and the α2-3 sub-helix. The approximate locations of the C and F pockets within the peptide binding groove are indicated with transparent letters. Residues 67-73 and 150-156 are labelled and coloured yellow and were used to measure the distance between α helices above the C pocket. Residues 74-85 and 138-149 were used similarly to measure the distance between α helices above the F pocket and are labelled and coloured orange. b) Two-dimensional frequency histograms of pocket distances from HLA-B*44:02 (top) and HLA-B*44:05 (bottom) across the second half of the simulation trajectory. C-pocket and F-pocket distances were measured, in angstroms, between the centre of mass of helix residues 67-73 and 150-156, and 74-85 and 138-149 respectively as in figure 1a. Approximate locations of energetic minima used to generate figure c are indicated by arrows in HLA-B*44:05. c) Representative cartoon structures for the identified C pocket distance minima in HLA-B*44:05 with Asn77 and Trp147 sidechains shown, for minima at 19 Å (top), 16 Å (centre) and 14 Å (lower). d) Structural depictions of the active displacement mechanism, showing reorientation of Tyr116 allowing competition for Trp147 hydrogen bond interactions (centre), leading to the hyper-closed conformation with the side chain of Trp147 reorientated outwards of the peptide binding groove.

We next sought to understand how this molecular rearrangement might control the tapasin-independent peptide editing capacity of HLA-B*44:05. In the peptide bound state, Asn77 shares a hydrogen bond with the amide bond linking the last two residues at the C-terminus of the peptide, Trp147 hydrogen bonds with the pΩ-1 carbonyl of the peptide, and Tyr116 sits at the base of the F pocket. Visualisation of the simulation trajectories revealed that in the closed conformation the Trp147 indole nitrogen was nearly perfectly aligned with the position of the peptide backbone amide in the peptide-loaded state (figure 2a). We hypothesised that in addition to its role as a specificity-determining sidechain in the F-pocket, Tyr116 may also function to compete with Asn77 for its hydrogen bonding partners. In the peptide-free closed conformation, this is Trp147, but in the peptide bound state it is the peptide backbone amide. Thus, we reasoned that Trp147 displacement out of the peptide binding groove, as seen in the hyper-closed conformation, serves as a proxy for peptide unbinding within our simulations. These simulations are consistent with a tapasin-independent peptide editing cycle in which peptide binding stability is iteratively sampled via the active displacement mechanism (figure 2b). Peptide exchange is most likely to occur in the “displaced” conformation, with active displacement sampling the aggregate strength of peptide-MHC-I interactions while the C-terminal amino acid of the peptide is displaced out of the peptide binding groove in HLA-B*44:05.

**Figure 2.**
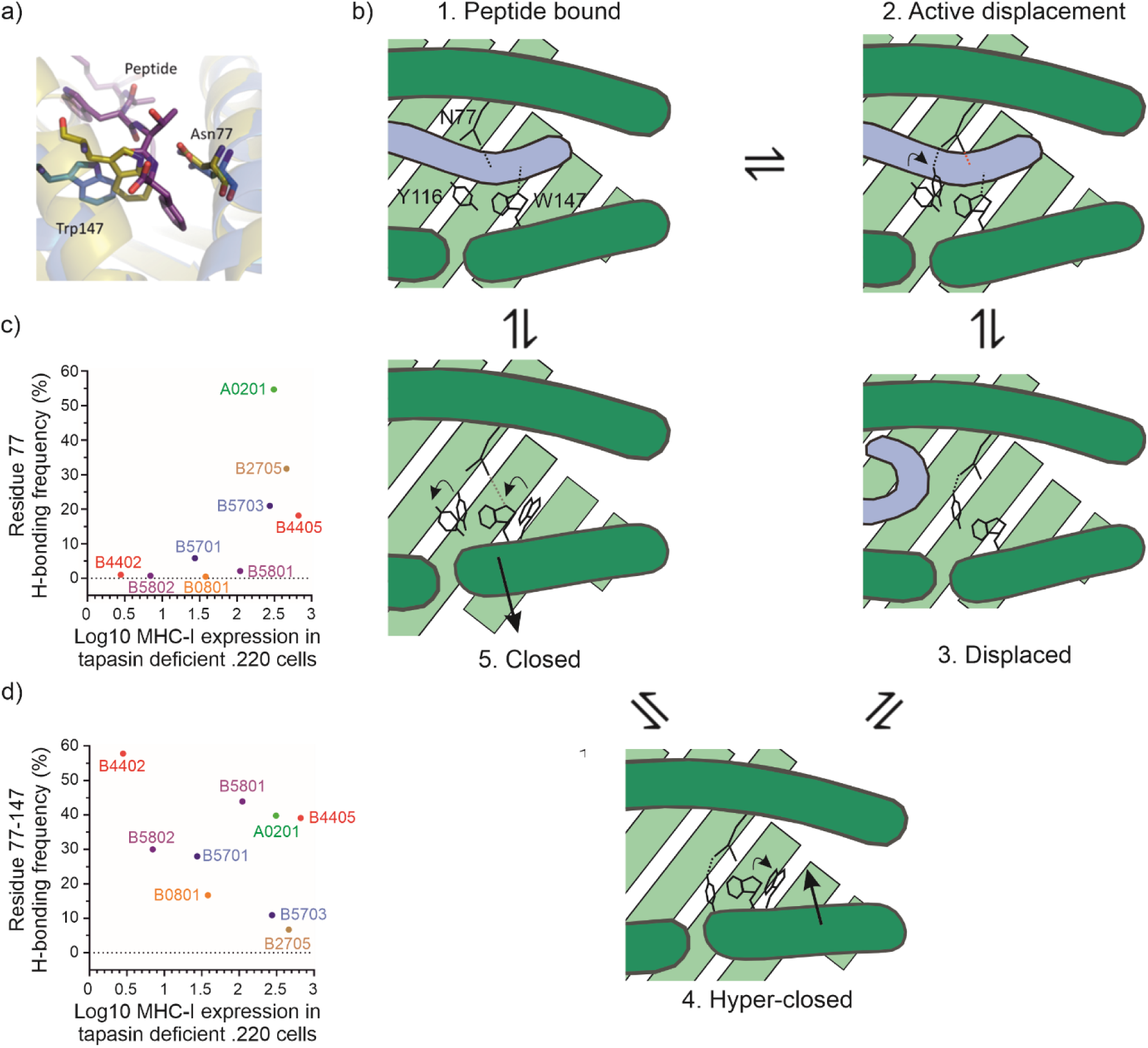
Dynamic hydrogen bonding involving residue 77 underpins independent peptide editing. a) Comparison of the positioning of the Asn77 and Trp147 side chains in HLA-B*44:05 in the presence or absence of peptide. The side chains are shown in blue in the peptide bound state, with peptide shown in purple, and overlaid with the orientations taken from a frame of the HLA-B*44:05 REST2 trajectory in the closed conformation (yellow). b) Graphical representation of the proposed catalytic independent peptide editing cycle, exemplified by HLA-B*44:05. In the peptide bound state (panel 1), multiple interactions hold the peptide within the peptide binding groove, including hydrogen bonds between the peptide backbone and MHC-I residues Asn77 and Trp147 (black dotted lines). Transient reorientation of the Tyr116 sidechain (panel 2) occurs constitutively (curved arrowhead) underpinning active displacement. This reorientation allows Tyr116 to transiently hydrogen bond with Asn77, which pulls Asn77 into an orientation that is incompatible with hydrogen bonding to the peptide backbone amide (red dashed line). Consequently, the C-terminus of the peptide may dissociate from the peptide binding groove (panel 3). In the absence of sufficient stabilising interactions with MHC-I, the peptide is completely released from the peptide binding groove (panel 4). Being unable to hydrogen bond with the peptide or with Asn77, the sidechain of Trp147 rotates (curved arrowhead) and MHC-I transitions to the “hyper-closed” conformation, with narrowing of the distance between alpha helices above the F pocket (thick inward pointing arrowhead) likely to preclude peptide binding. However, peptide binding is likely to be permitted once MHC-I reverts to a closed conformation (panel 5) with slight widening of the alpha helices (thick outward pointing arrowhead). For HLA-B*44:05, this is facilitated by reversion of Tyr116 to its original orientation (curved arrowhead), breaking the Asn77-Tyr116 hydrogen bond. Without the Asn77-Tyr116 bond, the sidechain of Trp147 resumes its former position (curved arrowhead) and shares a hydrogen bond with Asn77 (grey dashed line). Peptide binding to empty HLA-B*44:05 is likely to require the peptide to outcompete the Asn77-Trp147 hydrogen bond and is facilitated by the reorientation of Tyr116 disrupting the Asn77-Trp147 hydrogen bond. By contrast, peptide exchange is likely to occur in the displaced conformation (panel 3), with low affinity peptide dissociation being followed by breakage of the transient Asn77-Tyr116 hydrogen bond allowing binding of the replacement peptide. The molecular events underpinning active displacement, exemplified here by rotation of Tyr116, occur continuously allowing peptide affinity to be iteratively sampled. c) Graph showing the relationship between the frequency of residue 77 hydrogen bonding interactions and MHC-I cell surface expression levels previously measured in the absence of tapasin for the members of our MHC-I allotype panel (33). Potential residue 77 hydrogen bond frequency with polar sidechain groups was measured across REST2 MHC-I peptide-deficient simulations using a 3.5 Å distance interaction cut-off, excluding potential interaction with residues 74, 76, 78, 97, 123, 139-158 which are unlikely to interact with residue 77 in the presence of peptide bound state. MHC-I cell surface expression levels were previously measured in the absence of tapasin as in ref (33). MHC-I allotype pairs share coloured symbols (red: HLA-B*44:02/B*44:05, blue: HLA-B*57:01/B*57:03, purple: HLA-B*58:01/B*58:02), while single MHC-I allotypes (HLA-A*02:01, HLA-B*08:01, HLA-B*27:05) are denoted with unique coloured symbols. d) Graph showing the relationship between MHC-I cell surface expression levels previously measured in the absence of tapasin and the percentage of simulation frames in which Trp147 is hydrogen bonded to residue 77 for the members of our MHC-I allotype panel. The frequency of residue 77-147 hydrogen bond was measured using a 3.5 Å heavy atom cut-off distance across simulation times 50-100 ns. MHC-I allotypes are denoted with coloured symbols as in figure 2c.

To explore the generalisability of this observation further, we conducted additional REST2 simulations on a panel of MHC-I allotypes to determine the intrinsic capacity to disrupt the 77-147 hydrogen bond. Thus, we measured the frequency of residue 77 hydrogen bonding with nearby polar sidechain groups, excluding Trp147 and residues which were presumed to be sterically incapable of interacting with residue 77 or which made contacts to residue 77 in the peptide bound state. We found the hydrogen bond frequency was correlated with the MHC-I cell surface expression levels previously measured in the absence of tapasin (33) (figure 2c, supplementary figure 2 and tables 1+2). We observed that generally, allotypes with limited independent peptide editing capability (i.e. high tapasin-dependence and therefore low MHC-I expression in the absence of tapasin on the x-axis) did not have significant hydrogen bonding interactions involving residue 77. By contrast, MHC-I allotypes with greater independent peptide editing ability (i.e. lower tapasin-dependence, and higher MHC-I expression in the absence of tapasin) had more significant hydrogen bonding interactions involving residue 77 (figure 2c and supplementary figure 2). Notably, the more tapasin-dependent MHC-I allotype of each of the three allotype pairs had lower hydrogen bonding interactions involving residue 77.

An alternative possibility is that the Asn77-Trp147 hydrogen bond simply “locks” the peptide binding groove closed for tapasin-dependent HLA-B*44:02, but not for tapasin-independent HLA-B*44:05. We think this is unlikely because we found no correlation between the frequency of residue 77-Trp147 interactions and MHC-I cell surface expression levels previously measured in the absence of tapasin (figure 2d and table 1). Similarly, there was no correlation between peptide binding groove plasticity around the F and C-pockets and independent peptide editing ability across the panel of MHC-I allotypes, i.e. there was no consistent trends when tapasin-dependent and independent self-editing MHC-I allotype pairs were compared (supplementary figure 3).

**Table 1.** Correlation analyses of the relationships between the frequency of residue 77 hydrogen bonding to specific residues and MHC-I expression measured in the absence of tapasin. Spearman correlation analyses were performed to determine whether MHC-I cell surface expression measured in the absence of tapasin correlated with the frequency of hydrogen bonds involving residue 77 and specific residues. The correlation coefficient (r) is the fraction of variance that is shared between both variables. The p value represents the result of a test of the null hypothesis that the data were sampled from a population in which there is no correlation between the two variables.

|  | **Figure 2c**<br>**MHC-I expression in the absence of**<br>**tapasin vs the frequency of residue 77**<br>**hydrogen bonding to nearby polar**<br>**residues** | **Figure 2d**<br>**MHC-I expression in the absence**<br>**of tapasin vs the frequency of**<br>**residue 77 hydrogen bonding to**<br>**Trp147.** |
| --- | --- | --- |
| Spearman<br>r | 0.7333 | -0.3167 |
| P value | 0.0311 | 0.4101 |

### Experimental verification of the HLA-B*44:05 peptide editing mechanism

To test our hypothesis experimentally, we compared the independent peptide editing abilities of HLA-B*44:05 with a HLA-B*44:05-Y116F (Y116F) mutant. According to our hypothesis, the sterically conservative Phe116 would prevent hydrogen bonding with Asn77, thus disrupting the independent peptide editing ability of HLA-B*44:05. Accordingly, we did not observe any significant hydrogen bonds between Asn77 and nearby polar sidechain residues in Y116F REST2 simulations (table 2 and supplementary figure 2). We therefore produced recombinant HLA-B*44:05 and Y116F proteins and identified a panel of peptides that bound to both HLA-B*44:05 and Y116F proteins with very similar half-life (figure 3a, table 3). We conducted *in vitro* peptide exchange experiments comparing how each peptide competed against the fluorescent EEFGK^TAMRA^AFSF peptide for binding to HLA-B*44:05 or Y116F (figure 3b, table 3). Using this approach, we have previously demonstrated that when comparing MHC-I, reduced ability of a test peptide to compete with the fluorescent-labelled index peptide indicates a diminished ability of the MHC-I to undertake independent peptide editing (42). We observed that peptides with IC50s between 1-10 µM had a reduced ability to compete with EEFGK^TAMRA^AFSF for binding to B*44:05-Y116F compared to HLA-B*44:05 indicating that for these peptides Y116F was a poorer peptide editor. Peptides with IC50 values outside this range were less affected.

**Figure 3.**
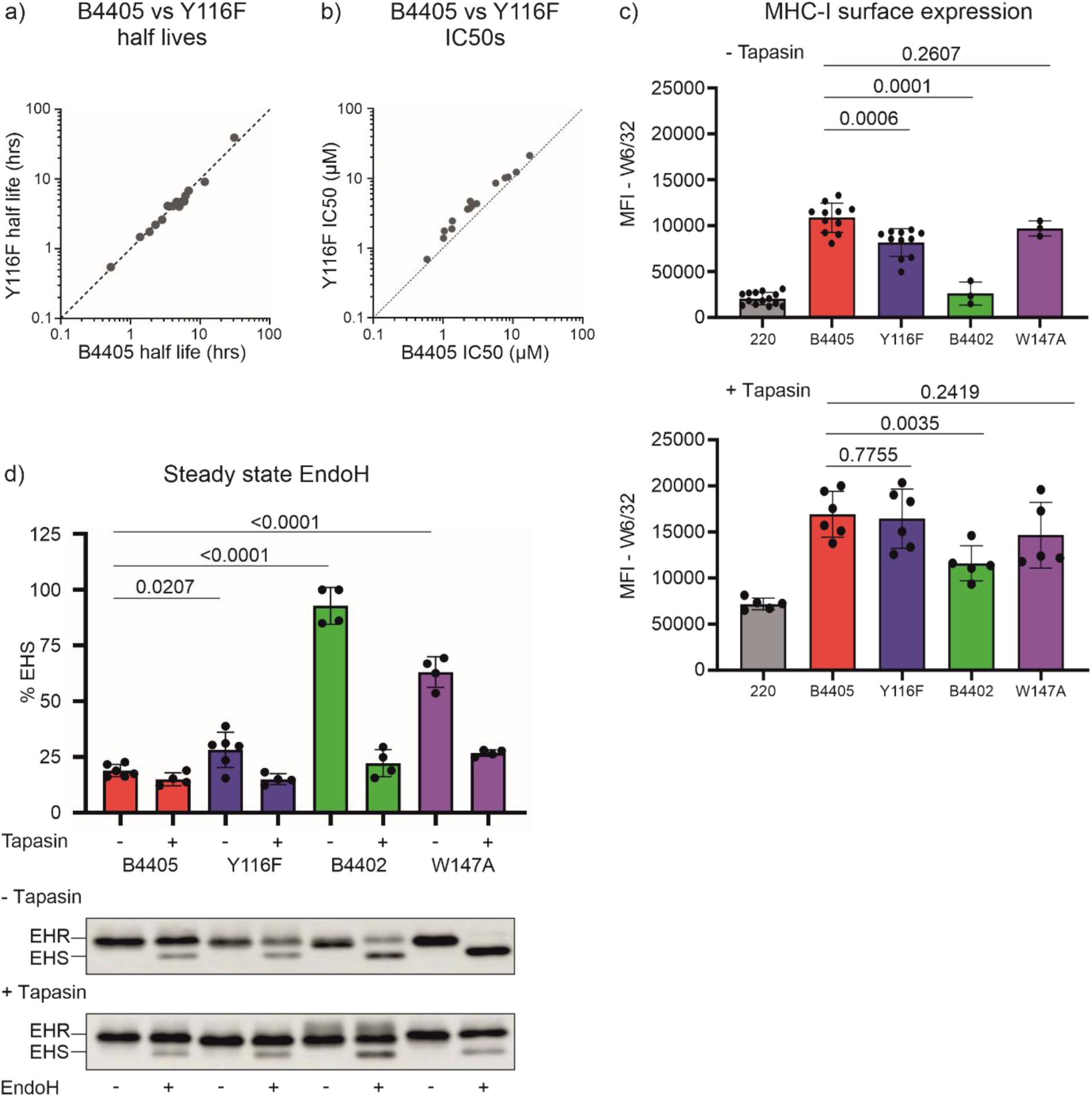
Experimental verification of the HLA-B*44:05 canonical active displacement mechanism. a) The half-lives of the complexes formed between the unlabeled peptides and HLA-B*44:05 or Y116F were indirectly measured as in ref (90). Each peptide was tested at least twice, with the mean half-lives reported in hours, with each peptide indicated by a grey dot. The mean half-lives of the peptide-HLA-B*44:05 complexes are shown on the x-axis, while the mean half-lives of the peptide-Y116F complexes are shown on the y-axis. b) Peptide competition experiments were conducted in which unlabelled peptides individually competed against EEFGK^TAMRA^AFSF peptide for binding to either HLA-B*44:05 or HLA-B*44:05-Y116F. Binding of EEFGK^TAMRA^AFSF peptide was measured by fluorescence polarisation and IC50 values calculated for each peptide (IC50 value is shown in µM units, with high affinity peptides having low µM values). Each peptide was tested at least twice, and the mean of the replicate experiments is reported. Peptides with equal ability to compete for binding to wild-type and Y116F HLA-B*44:05 molecules will fall along the diagonal dashed line. c) Bar graphs showing the cell surface expression levels of the indicated MHC-I molecules expressed in tapasin-deficient 721.220 cells (no tapasin, upper) or tapasin-reconstituted 721.220-tapasin cells (+ tapasin, lower) following staining with W6/32 antibody and flow cytometry analysis. The bar height represents the mean expression levels measured in multiple independent experiments, with dots representing the results of individual experiments, and vertical lines depicting the standard deviation observed between experiments. Differences in MHC-I expression are marked with horizontal lines and their p value. d) Bar graph depicting the percentage of MHC-I molecules that were sensitive to digestion with endoglycosidase H. Lysates were prepared from transfectants expressing the indicated MHC-I molecules in the absence (- on x axis) or presence of tapasin (+ on x axis). The lysates were digested with endoglycosidase H, or mock digested, and separated by SDS PAGE, western blotted, and heavy chain bands detected using HC10 antibody. The bar height represents the mean percentage of MHC-I molecules that were endoH sensitive measured in multiple independent experiments, with dots representing the results of individual experiments, and vertical lines depicting the standard deviation observed between experiments. Differences in MHC-I expression are marked with horizontal lines and their p value. The lower panels show representative images of the indicated MHC-I heavy chain bands expressed in tapasin-deficient 721.220 cells (- tapasin, upper) or tapasin-reconstituted 721.220-tapasin cells (+ tapasin, lower) that were digested with endoglycosidase H, or mock digested (indicated with + or –). EHR and EHS denote immunoprecipitated MHC-I molecules that are resistant (EHR) or sensitive (EHS) to digestion with endoglycosidase H.

**Table 2.** Summary of residue 77 hydrogen bond statistics as measured by REST2 simulations. Residue 77 hydrogen bond frequency refers to percentage of simulation frames in which residue 77 shares a hydrogen bond to a compatible polar atom.

| **MHC-I**<br>**allotype** | **Mutation** | **Residue 77**<br>**hydrogen**<br>**bond**<br>**Frequency**<br>**(%)** |
| --- | --- | --- |
| B*44:02 | None | 1.03 |
| B*44:05 | None | 18.16 |
| B*44:05 | Y116F | 0.38 |
| B*57:01 | None | 5.80 |
| B*57:03 | None | 20.99 |
| B*58:01 | None | 2.11 |
| B*58:02 | None | 0.72 |
| B*08:01 | None | 0.47 |
| B*27:05 | None | 31.73 |
| A*02:01 | None | 54.72 |
| C*05:01 | None | 11.86 |
| C*05:01 | N77S | 0.00 |
| C*05:01 | K80N | 4.15 |
| C*08:02 | None | 0.17 |

**Table 3.**
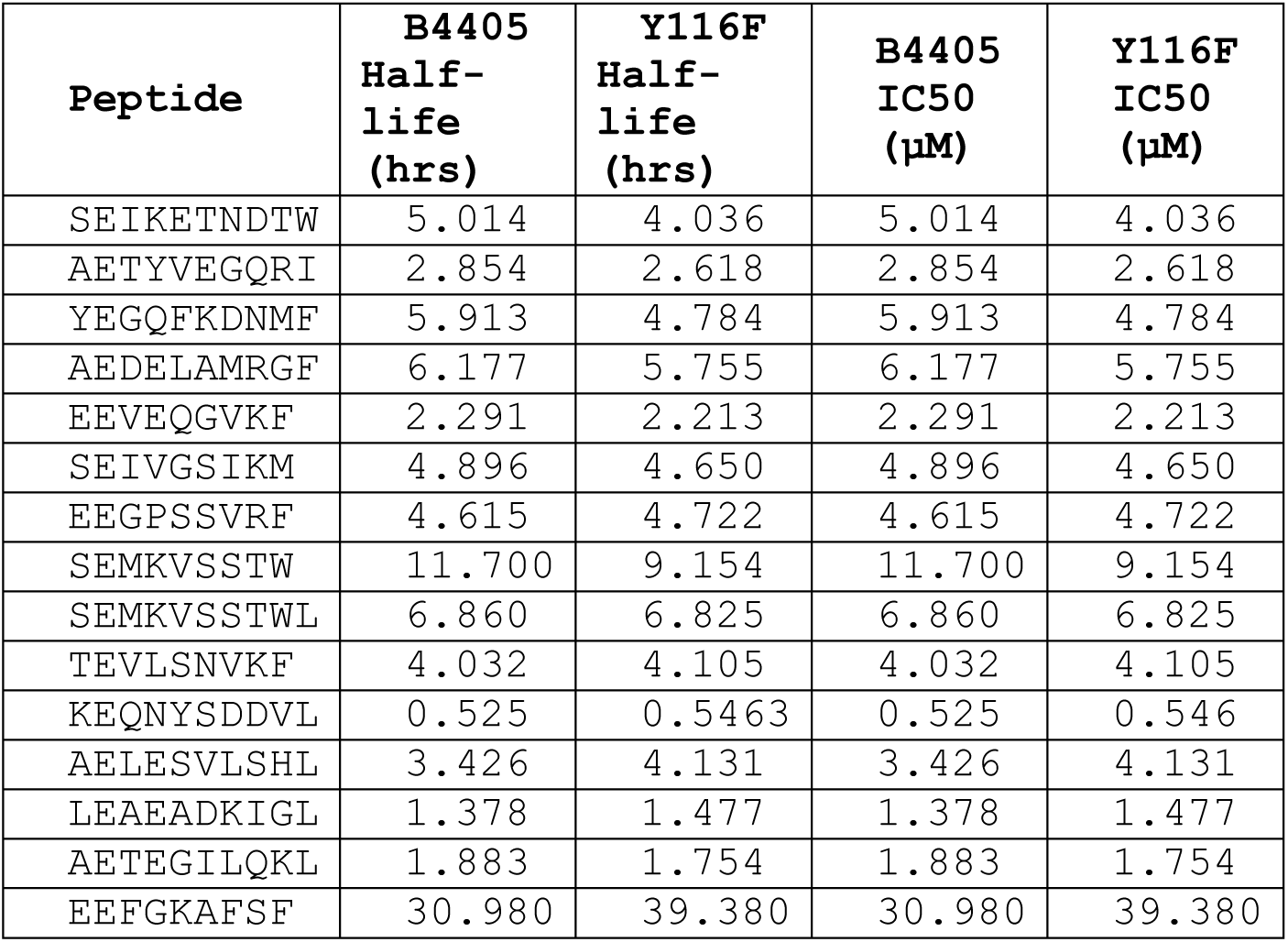
Half-lives and IC50 values measured for the indicated peptides competing against EEFGK^TAMRA^AFSF for binding to the HLA-B*44:05 or HLA-B*44:05-Y116F.

We next expressed the Y116F mutant in the tapasin-deficient 721.220 cell line, and its tapasin-reconstituted counterpart, and compared its peptide loading characteristics with those of HLA-B*44:02 or HLA-B*44:05. We also included in these experiments the HLA-B*44:05-W147A (W147A) mutant, which has level of tapasin-dependence that is intermediate between HLA-B*44:02 and HLA-B*44:05 (42, 51). As previously observed, in the absence of tapasin there was low surface expression of HLA-B*44:02, and high surface expression of HLA-B*44:05 (figure 3c, upper) (22, 33, 35, 53), thus defining the dynamic range of MHC-I expression in the absence of tapasin. The expression level of Y116F was significantly decreased compared with HLA-B*44:05, which was restored upon co-expression of tapasin. Thus, Y116F appears to reduce tapasin-independent peptide editing of HLA-B*44:05 in live cells. The Y116F molecules that egressed to the cell surface in the absence of tapasin are nevertheless stable indicating that the mutation did not enable HLA-B*44:05 to breech a quality control checkpoint upstream of peptide editing (Supplementary figure 4).

Seeking further evidence that the Y116F mutation impedes independent peptide editing, we digested lysates with endoglycosidase H (endoH) and compared the proportion of MHC-I molecules that were susceptible to digestion following SDS PAGE, western blotting and detection with HC10 antibody (figure 3d). Nascent MHC-I molecules acquire resistance to endoH when their peptide cargo permits their progression to the medial Golgi where their N-linked glycans are modified to become resistant to endoH (22, 54). In tapasin-deficient cells most HLA-B*44:02 and W147A molecules were sensitive to endoH, while most HLA-B*44:05 molecules were resistant to endoH, confirming earlier observations that tapasin-independent peptide editing by HLA-B*44:05 permits egress to the cell surface. Consistent with the cell surface expression levels, we observed a statistically significant increase in the proportion of Y116F molecules that were sensitive to endoH compared with HLA-B*44:05, though the effect was not so great as for HLA-B*44:02 or B*44:05-W147A. This difference was not apparent for any of the variants in tapasin-proficient cells, indicating that the effect was most likely due to tapasin-independent peptide editing.

Cumulatively, these experiments showed the Y116F mutation had a clear effect on individual peptides competing for binding to MHC-I and in cells, and that there was a reduction in Y116F cell surface expression levels that correlated with an increased proportion of unloaded or poorly loaded Y116F molecules that remained sensitive to endoglycosidase H. Comparison of HLA-B*44:05 and Y116F therefore supports our hypothesis that active displacement, involving dynamic hydrogen bonding involving Asn77, Tyr116 and Trp147 underpins the ability of HLA-B*44:05 to sample peptides for presentation without the assistance of tapasin.

### HLA-C independent peptide editing

We next sought to determine whether peptide cargo editing via active displacement may explain instances where related tapasin-independent MHC-I allotypes with the same specificity-determining F-pocket demonstrate a degree of fine-specificity in their binding preference.

HLA-C*05:01 and HLA-C*08:02 differ only at two positions (HLA-C*05:01: Asn77+Lys80, HLA-C*08:02: Ser77+Asn80), share similar peptide binding specificities (55, 56) and are both relatively tapasin-independent (33). However, the immunogenic KRAS-G12D neoantigen, GADGVGKSA was shown to be exclusively HLA-C*08:02 restricted and showed no T cell reactivity in the context of HLA-C*05:01 (57–60). The GADGVGKSA epitope is predicted to bind both allotypes with low affinity, yet in experiments this epitope bound and stabilised HLA-C*08:02 and not HLA-C*05:01 (57, 58), likely explaining the T cell restriction to HLA-C*08:02. However, structural studies could not explain why the GADGVGKSA epitope bound to HLA-C*08:02 and not HLA-C*05:01 (57). The crystal structure of GADGVGKSA in HLA-C*08:02 revealed canonical interactions with the peptide binding groove, with an empty cavity in the bottom of the F pocket, typically occupied by longer p9 sidechains (57–59). HLA-C*05:01 has an identical F pocket but was only stabilised by a p9 Leu variant peptide (GADGVGKSL) and not GADGVGKSA (57, 58). We hypothesised that active displacement might remove GADGVGKSA from the HLA-C*05:01 peptide binding groove, but not HLA-C*08:02.

We therefore performed REST2 simulations with HLA-C*05:01 and HLA-C*08:02. We found no differences in C or F pocket distances between either allotype indicating that global peptide binding groove dynamics do not underpin differences in the presentation of GADGVGKSA (figure 4a). However, we observed significant differences in hydrogen bonding dynamics involving residue 77 in each allotype: Ser77 in HLA-C*08:02 did not have any significant interactions, while Asn77 in HLA-C*05:01 had substantially higher hydrogen bonding frequency involving residue 77 and nearby polar sidechain residues besides Trp147 (figures 4b+4c, supplementary figures 2 and 5, and table 2). For HLA-C*05:01, we observed an alternative hydrogen bonding interaction in which Asn77 hydrogen bonded to Lys80 with a frequency of 3.4% in the simulations, reminiscent of interactions between Asn77 and Tyr116 in HLA-B*44:05 (figure 4b indicated by an orange arrowhead, with an enlarged view in supplementary figure 5, figure 4d right panel and supplementary figure 2). We hypothesised residue 80 facilitates active displacement in HLA-C*05:01 by disrupting the hydrogen bonding interaction between residues 77-147 in an analogous fashion to that we propose for HLA-B*44:05 (the proposed HLA-C independent editing cycle is depicted in figure 4e). For HLA-C*05:01, we also observed a hydrogen bonding interaction between Asn77 and Asn74 (figure 4b, indicated with a red arrowhead and supplementary figure 5). As the Asn77-Asn74 hydrogen bond is present in the available crystal structures of HLA-C*05:01 this interaction is unlikely to mediate active displacement and may instead stabilise the Asn77-peptide bond.

**Figure 4.**
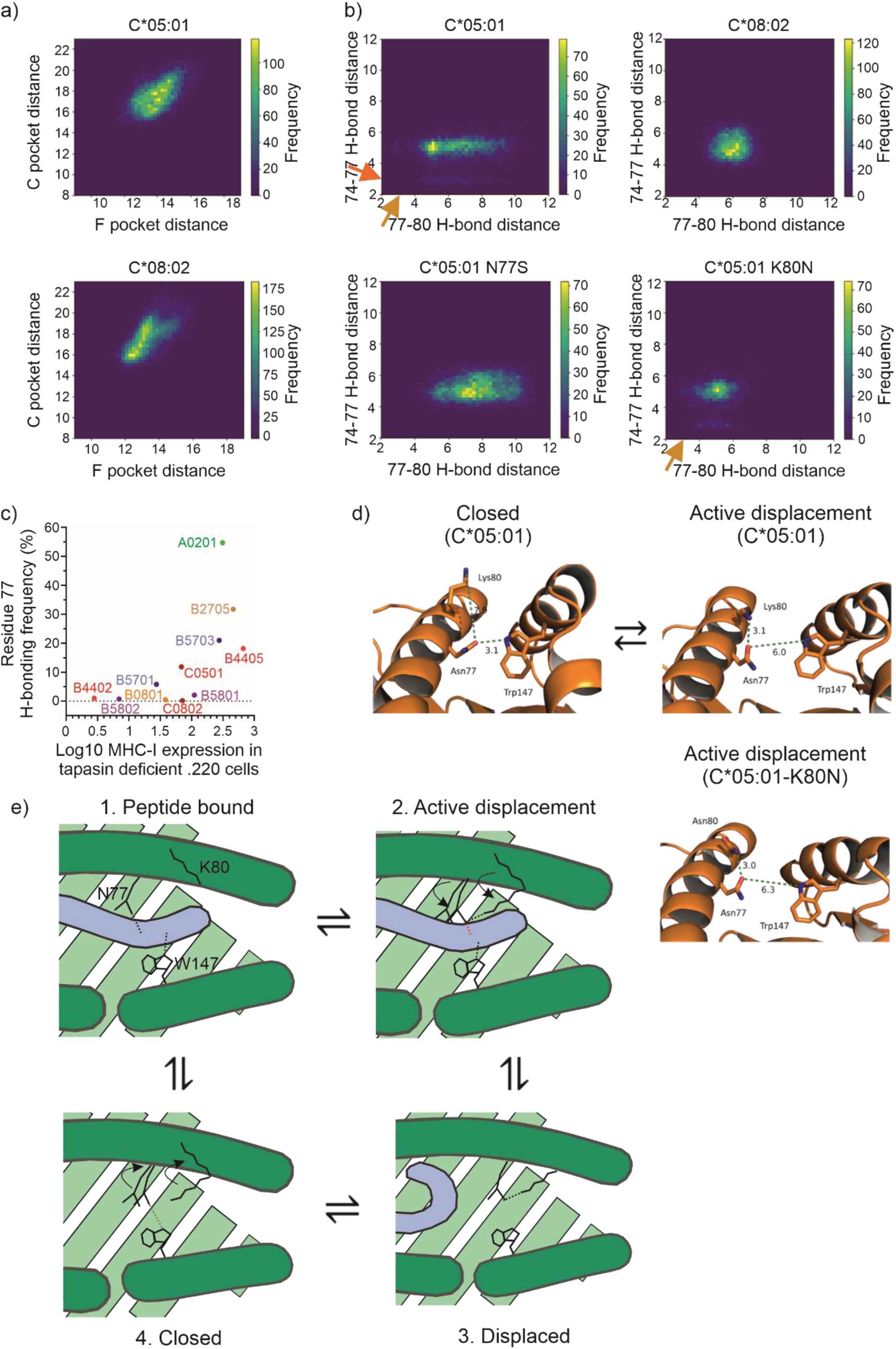
Identification of the HLA-C active displacement mechanism. a) Two-dimensional frequency histograms of pocket distances, in angstroms, from HLA-C*05:01 (top) and HLA-C*08:02 (bottom) across the second half of REST2 simulation trajectories. Pocket distances were calculated as in figure 1b. b) Two-dimensional frequency histograms of the minimum distance between residues 77 and 80 (x-axis) or 77 and 74 (y-axis) were used to identify hydrogen bonds at approximately 3-3.5 angstroms. Relevant potential hydrogen bond interactions discussed in the text are indicated with arrowheads. c) Graph showing the relationship between MHC-I cell surface expression levels previously measured in the absence of tapasin and the frequency of residue 77 hydrogen bonding interactions of HLA-C*05:01 and HLA-C*08:02 in the context of our MHC-I allotype panel as described in figure 2c. d) Structural depictions of the HLA-C active displacement mechanism. Simulation snapshots compare the closed conformation of HLA-C*05:01 (left) with the proposed active displacement mediating conformations of HLA-C*05:01 (right). The reorientation of residue 80 strains the Asn77-Trp147 bond that acts as a surrogate for the peptide backbone as described for HLA-B*44:05. The lower panel depicts how the Asn80 side chain of C*05:01-K80N is also able to hydrogen bond with Asn77 to bring about active displacement. e) Graphical representation of the proposed HLA-C catalytic independent peptide editing cycle, exemplified by HLA-C*05:01. In the peptide bound state (panel 1), multiple interactions hold the peptide within the peptide binding groove, including hydrogen bonds between the peptide backbone and MHC-I residues Asn77 and Trp147 (black dotted lines). Transient reorientation of the Lys80 sidechain (panel 2) occurs constitutively (curved arrowhead) underpinning HLA-C active displacement. This reorientation allows Lys80 to transiently hydrogen bond with Asn77, which pulls Asn77 (curved arrowhead) into an orientation that is incompatible with hydrogen bonding to the peptide backbone amide (red dashed line). Consequently, the C-terminus of the peptide may dissociate from the peptide binding groove (panel 3). In the absence of sufficient stabilising interactions with MHC-I, the peptide is completely released from the peptide binding groove (panel 4). Reversion of Lys80 to its original orientation leads to the repositioning of Asn77 into its original position (curved arrowheads) where it shares a hydrogen bond with Trp147 (grey dashed line). Peptide binding to empty HLA-C*05:01 is likely to require the peptide to outcompete the Asn77-Trp147 hydrogen bond and is facilitated by the transient reorientation of Lys80 disrupting the Asn77-Trp147 hydrogen bond. By contrast, peptide exchange is likely to occur in the displaced conformation (panel 3), with low affinity peptide dissociation being followed by breakage of the transient Asn77-Lys80 hydrogen bond allowing binding of the replacement peptide. The molecular events underpinning HLA-C active displacement, exemplified here by rotation of Lys80, occur continuously allowing peptide affinity to be iteratively sampled.

To test the hypothesis that residues 77 and 80 in HLA-C*05:01 are involved in active displacement, we examined single point mutants, HLA-C*05:01-N77S (C*05:01-N77S) and HLA-C*05:01-K80N (C*05:01-K80N). Sim et al previously showed that GADGVGKSA specific TCR could detect its cognate epitope loaded on C*05:01-N77S, but not C*05:01-K80N (57). We found that in C*05:01-N77S, residue 77 showed no hydrogen bonding to residue 74 or 80, likely due to the shorter serine sidechain being unable to reach potential hydrogen bonding partners (figure 4b, supplementary figure 5, and table 2). By contrast in the simulations of C*05:01-K80N, we observed the hydrogen bond interaction between residues 77-80 with a frequency of 4.0% (figure 4b, indicated with an orange arrowhead, supplementary figure 5, and table 2). Therefore, our simulations indicate that in HLA-C*05:01 and C*05:01-K80N, residue 80 disrupts the Asn77-Trp147 hydrogen bond (figure 4d right panels), which serves as a proxy for the Asn77-peptide backbone interaction in our simulations. This disruption is likely to lead to the release of the low affinity peptides, such as GADGVGKSA, via active displacement (figure 4e). By contrast in HLA-C*08:02 and C*05:01-N77S, residue 80 is unable to disrupt the Ser77-Trp147 hydrogen bond and consequently GADGVGKSA is not displaced from the peptide binding groove thus allowing T cell detection.

### Experimental verification of the HLA-C peptide editing mechanism

We next tested our hypothesis using *in vitro* peptide competition experiments. We first designed a fluorescently labelled tracer peptide (SAEPK^TAMRA^PLQL) based on the HLA-C*05:01-SAEPKPLQL crystal structure (61). When loaded on HLA-C*08:02, HLA-C*05:01, C*05:01-N77S and C*05:01-K80N, unlabelled SAEPKPLQL competed with SAEPK^TAMRA^PLQL peptide for binding with very similar IC50s (figure 5a and table 4). In sharp contrast, the KRAS-G12D neoantigen GADGVGKSA displayed considerably different IC50s to the HLA-C variants with single mutations at positions 77 or 80. GADGVGKSA competed easily against the SAEPK^TAMRA^PLQL peptide for binding to HLA-C*08:02 and C*05:01-N77S, but showed almost no ability to compete with the tracer peptide for binding to HLA-C*05:01 or C*05:01-K80N (figure 5b, table 4). This is consistent with a mechanism of active displacement in which the dynamic internal hydrogen bond networking permitted by HLA-C*05:01 and C*05:01-K80N displaces GADGVGKSA resulting in poor competition for the tracer peptide.

**Figure 5.**
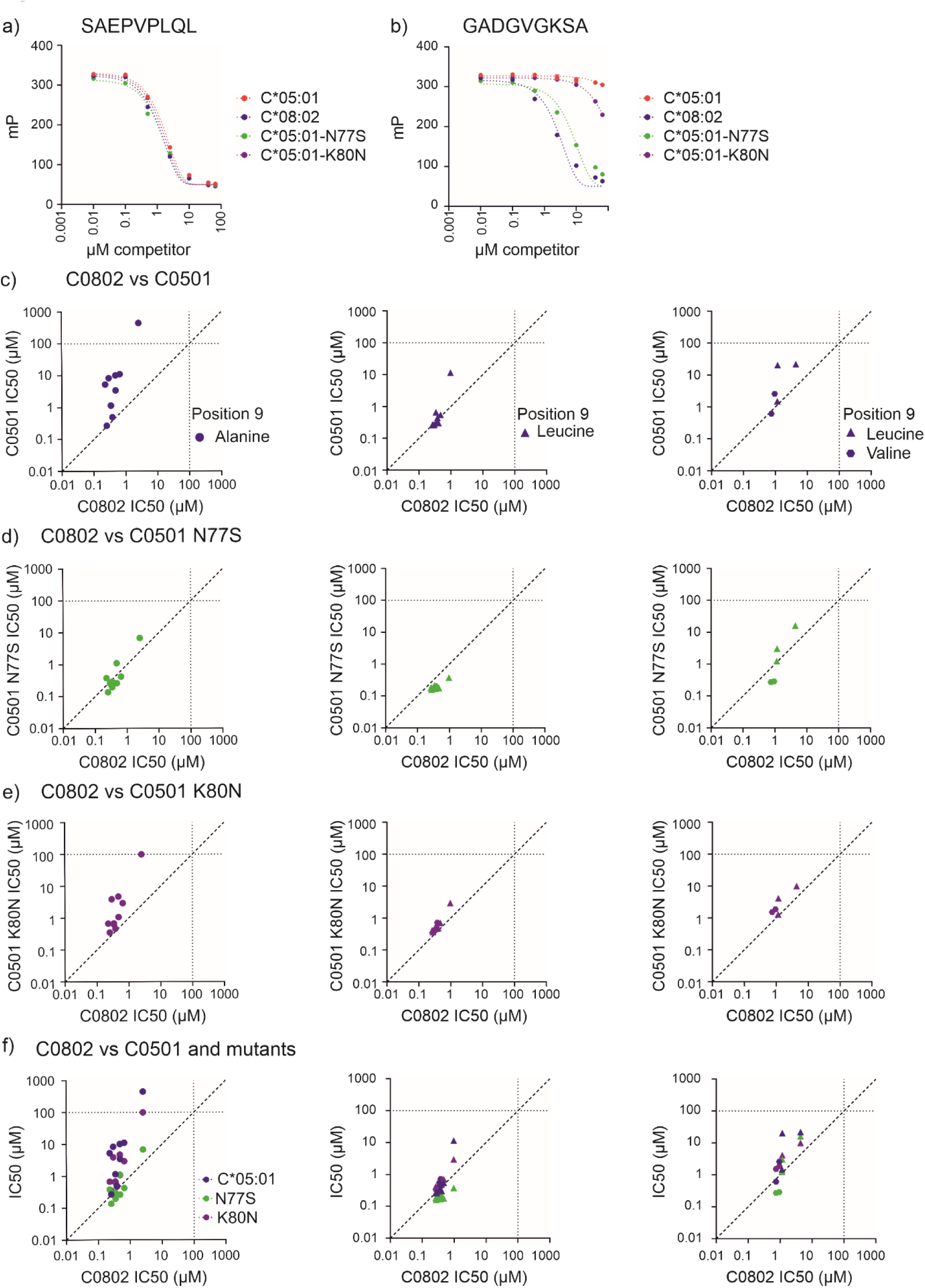
Experimental verification of the HLA-C active displacement mechanism. a) An *in vitro* peptide competition experiment was conducted in which unlabelled SAEPKPLQL peptide competed against SAEPK^TAMRA^PLQL peptide for binding to either HLA-C*05:01, HLA-C*08:02, C*05:01-N77S or C*05:01-K80N. Binding of SAEPK^TAMRA^PLQL peptide was measured by fluorescence polarization and is reported in mP as described in the methods. Unbound SAEPK^TAMRA^PLQL has a polarisation of 50 mP, while HLA-C bound SAEPK^TAMRA^PLQL has a polarisation of ∼300 mP. b) An *in vitro* peptide competition experiment was conducted in which unlabelled GADGVGKSA peptide competed against SAEPK^TAMRA^PLQL peptide for binding to either HLA-C*05:01, HLA-C*08:02, C*05:01-N77S or C*05:01-K80N as in figure 5a. c) Peptide competition experiments were conducted in which unlabelled peptides individually competed against SAEPK^TAMRA^PLQL peptide for binding to either HLA-C*05:01 or HLA-C*08:02. IC50 values were calculated for each peptide as in figures 3a+b. The panels compare IC50 values measured for peptides with alanine at position 9 (p9A, cohort 1, left), peptides with leucine at position 9 (p9L, cohort 2, centre), or peptides from cohort 3 (others, right) competing for binding to either HLA-C*05:01 (y axes), or HLA-C*08:02 (x axes). Each peptide was tested twice, and the mean of the replicate experiments is reported. Peptides with equal ability to compete for binding against the TAMRA-labelled peptide to both MHC-I molecules will fall along the diagonal dashed line. Faint dashed lines indicate 100 µM concentrations to facilitate visualization of data, and each peptide is denoted by as in the key. d) Peptide competition experiments were conducted as in figure 5c with peptides competing for binding to HLA-C*05:01 N77S or HLA-C*08:02. e) Peptide competition experiments were conducted as in figure 5c with peptides competing for binding to HLA-C*05:01 K80N or HLA-C*08:02. f) Plots summarising how each cohort of peptides competed for binding to the indicated HLA-C*05:01 molecules (y axes) in relation to HLA-C*08:02 (x axes).

**Table 4.** IC50 values measured for the indicated peptides competing against SAEPK^TAMRA^PLQL for binding to the HLA-C*05:01, HLA-C*08:02, C*05:01-N77S or C*05:01-K80N.

| Cohort | Peptide | C*08:02 | C*05:01 | C*05:01-N77S | C*05:01-K80N |
| --- | --- | --- | --- | --- | --- |
| 1 | GADGVGKSA | 2.549 | 448.700 | 6.867 | 99.920 |
|  | YADEKGLEA | 0.348 | 1.155 | 0.197 | 0.630 |
|  | FADAHSNLA | 0.383 | 0.500 | 0.267 | 0.468 |
|  | FADQENQVA | 0.255 | 0.270 | 0.138 | 0.354 |
|  | LSDDPTVSA | 0.295 | 8.410 | 0.258 | 3.927 |
|  | VSDTPSITA | 0.484 | 3.463 | 0.265 | 1.080 |
|  | AADLAHITA | 0.654 | 11.180 | 0.427 | 2.963 |
|  | VADGLPLAA | 0.477 | 10.250 | 1.121 | 4.737 |
|  | LADELINAA | 0.229 | 5.288 | 0.382 | 0.675 |
|  | LTDTPDTTA | 0.345 | 1.163 | 0.295 | 0.684 |
| 2 | GADGVGKSL | 0.969 | 11.500 | 0.378 | 2.964 |
|  | YADEKGLEL | 0.263 | 0.264 | 0.181 | 0.415 |
|  | FADAHSNLL | 0.411 | 0.301 | 0.190 | 0.486 |
|  | FADQENQVL | 0.276 | 0.269 | 0.159 | 0.371 |
|  | LSDDPTVSL | 0.281 | 0.260 | 0.181 | 0.364 |
|  | VSDTPSITL | 0.369 | 0.338 | 0.169 | 0.467 |
|  | AADLAHITL | 0.476 | 0.545 | 0.178 | 0.705 |
|  | VADGLPLAL | 0.380 | 0.438 | 0.204 | 0.721 |
|  | LADELINAL | 0.340 | 0.657 | 0.218 | 0.557 |
|  | LTDTPDTTL | 0.301 | 0.269 | 0.168 | 0.378 |
| 3 | SAEPVPLQL | 1.150 | 1.495 | 1.249 | 1.292 |
|  | FANMGIALTL | 4.376 | 21.920 | 16.220 | 10.000 |
|  | KTDGTGVHATL | 1.168 | 20.370 | 3.062 | 4.159 |
|  | IIDKSGIPV | 0.945 | 2.587 | 0.289 | 1.876 |
|  | IIDKSGSEV | 0.747 | 0.612 | 0.272 | 1.533 |

To extend this finding we measured the ability of three peptide cohorts to compete for binding to HLA-C*05:01, HLA-C*08:02, C*05:01-N77S or C*05:01-K80N (table 5). The first cohort were peptides that had been eluted from HLA-C*08:02 containing C-terminal position 9 Ala (56). Of note, peptides with C-terminal alanine were found in the immunopeptidomes of HLA-C*08:02 and HLA-C*05:01 at a frequency of 1.5% and 0.13% respectively (57). We reasoned these peptides may also be susceptible to active displacement in the context of HLA-C*05:01 and C*05:01-K80N. The second cohort were otherwise identical in sequence to the first with leucine at position 9, rather than a sub-optimal alanine. The third cohort consisted of five peptides, all of which have preferred C-terminal residues and included the unlabelled version of the fluorescent tracer peptide, two variants of the IIDKSGSTV peptide differentially recognised by killer immunoglobulin-like receptors (62), and two neoantigens predicted to bind both HLA-C*05:01 and HLA-C*08:02 (63–65).

**Table 5.**
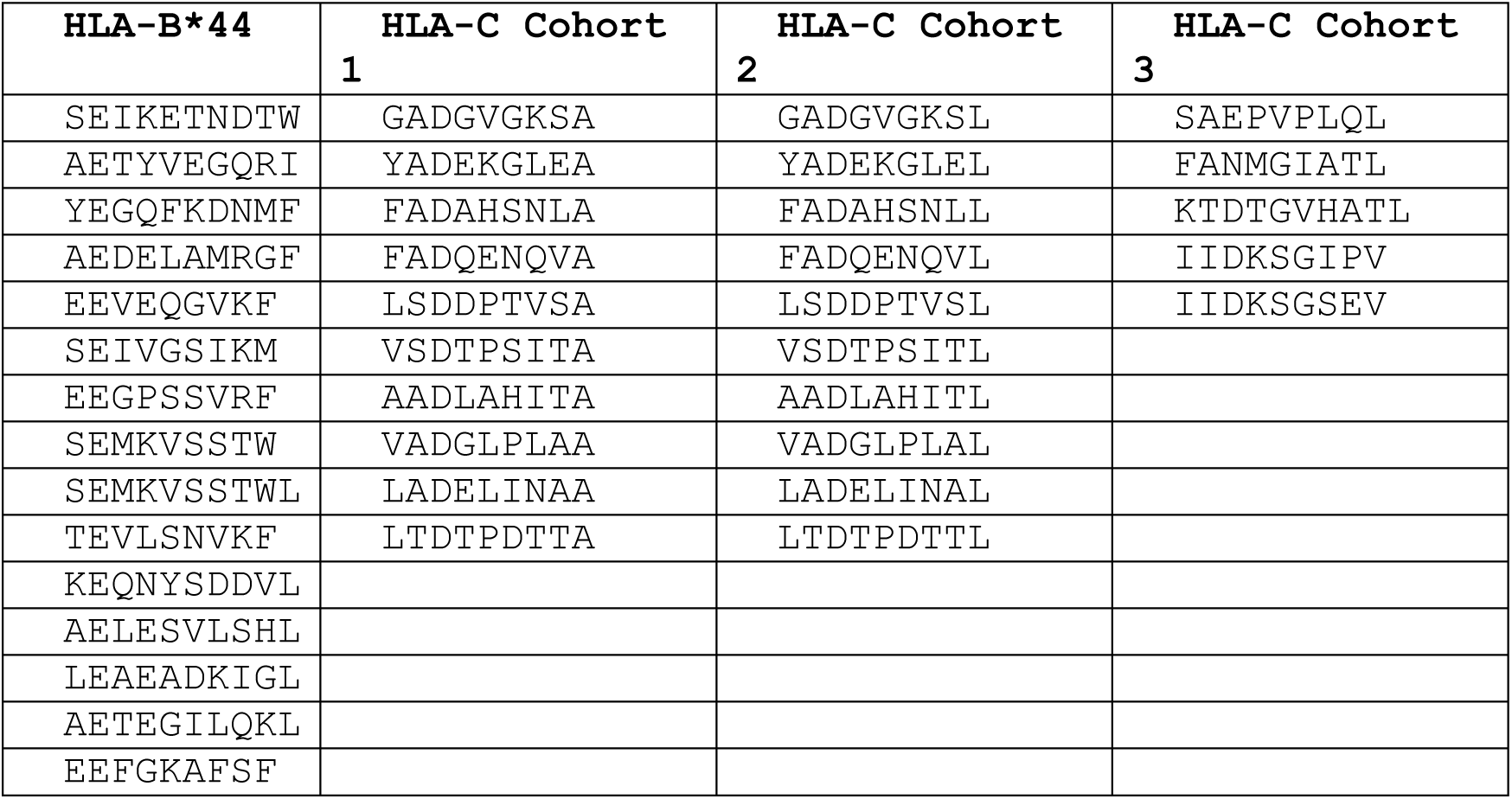
HLA-B*44 and HLA-C competitor peptide sequences.

Most peptides in cohort 1 were poor competitors for binding to HLA-C*05:01 compared to HLA-C*08:02, indicated by higher IC50 values (figure 5c, left panel, table 4). However, this was not the case for the position 9 leucine equivalent peptides in cohort 2, where, apart from GADGVGKSL, the peptides had a similarly strong ability to compete for binding to either HLA-C*05:01 or HLA-C*08:02 (figure 5c, centre panel, table 4). These results are consistent with HLA-C*05:01, but not HLA-C*08:02, having an intrinsic peptide editing mechanism that displaces peptides with suboptimal position 9 anchors from the peptide binding groove. Two of the 10 peptides within cohort 1, FADAHSNLA and FADQENQVA, had a similarly strong ability to compete for binding to HLA-C*05:01 as to HLA-C*08:02 (table 4). Interestingly, these were the only peptides with phenylalanine at their N-terminus. Phenylalanine is the most common p1 residue found in the C*05:01 immunopeptidome (56) and thus may optimally fill the A pocket potentially counteracting the active displacement mechanism despite a suboptimal P9 anchor.

We found that most peptides in cohort 3 were poorer competitors for binding to HLA-C*08:02 than the peptides from the other cohorts (figure 5c and table 4). Despite having preferred C-terminal residues, the peptides FANMGIATL, KTDTGVHATL and IIDKSGIPV were substantially poorer competitors for binding to HLA-C*05:01 than they were for HLA-C*08:02 (table 4). This was also true for GADGVGKSL, the lowest affinity competitor in cohort 2, suggesting that the active displacement mechanism could remove low affinity peptides from the peptide binding groove despite these peptides having optimal p9 anchor residues. The active displacement mechanism outlined above to describe tapasin-independent repertoire editing by HLA-B*44:05 (figure 2b) can therefore be extended to include the example of “repertoire refinement” seen for HLA-C*08:02, as outlined in figure 4e. By analogy it follows that any weaker binding peptide, regardless of C-terminal anchor should be susceptible to editing by C*08:02, as exemplified by GADGVGKSL, FANMGIATL, KTDTGVHATL and IIDKSGIPV.

In contrast to HLA-C*05:01, we found that all three cohorts of peptides competed for binding to C0501-N77S in a similar fashion as they did to HLA-C*08:02 (figures 5d and 5f, and table 4). However, peptide competition for binding to C*05:01-K80N, was similar to HLA-C*05:01 (figures 5e and 5f, and table 4). Thus, active displacement determined by sequence at positions 77 and 80 explains why HLA-C*05:01 and C*05:01-K80N could not stimulate KRAS neoepitope GADGVGKSA specific Jurkat cells (57). These data support a model where active displacement can account for tapasin-independent peptide-editing with ramifications including T cell recognition of cancer antigens.

## Discussion

The presentation of peptides by MHC-I molecules is essential for T cell medicated adaptive immune responses. The highly polymorphic and diverse nature of MHC-I molecules influences several functional properties. In humans, MHC-I allotypes have been shown to differ in: MHC-I cell surface expression levels (43–45); immunopeptidome diversity (41, 42, 44, 45); and dependence upon tapasin or TAPBPR for peptide acquisition (33, 34, 66, 67). Recently, the dependence upon tapasin for cell surface expression of HLA-A and HLA-B allotypes was shown to inversely correlate with the number of peptides derived from HIV eliciting immune responses (33). Similarly, MHC-I immunopeptidome diversity and cell surface expression levels were shown to be inversely correlated, and that this relationship correlated with resistance to certain infectious pathogens in chickens and humans (36, 43). Chappell et al extended these findings by noting that tapasin-dependence correlated with MHC-I cell surface expression levels (35, 43). Collectively, these studies suggest that tapasin-dependence underpins MHC-I cell surface expression level and immunopeptidome diversity, and that these properties synergistically influence immune responses. The expression levels of tapasin, MHC-I, and other components of the peptide-loading complex (PLC) can vary dynamically, being upregulated by cytokine signalling to enhance antigen presentation or downregulated by pathogens (39, 40, 68, 69) and cancer (70, 71) to escape detection. Divergence in dependence upon tapasin for MHC-I cell surface expression may have been evolutionarily selected to counter the threat of disruption to tapasin-mediated peptide editing. However, a universal explanation for why some MHC-I allotypes can self-edit while others rely on tapasin remains elusive. Here, we provide a molecular model, which we have termed active displacement, accounting for the ability of some MHC-I allotypes to independently edit their peptide repertoire.

By conducting enhanced sampling molecular dynamic simulations on a panel of MHC-I allotypes, we identified two related atomistic mechanisms of independent peptide editing by MHC-I involving dynamic hydrogen bonding of Asn77. Importantly, both culminate in the transient disruption of the hydrogen bond interaction between Asn77 of the MHC-I and the amide bond linking pΩ and pΩ-1 of the peptide. For HLA-B*44:05, Tyr116 is the effector sidechain (figure 2b) while Lys80 is the effector for HLA-C*05:01 (figure 4e). Reorientation of these effector sidechains allows the effectors to transiently share a hydrogen bond with Asn77, which pulls Asn77 into an orientation that is incompatible with hydrogen bonding to the peptide backbone amide. The computationally intensive nature of enhanced sampling molecular dynamic simulations prohibited us from modelling the atom-level events that may lie downstream of these molecular rearrangements that lead to peptide dissociation. As depicted in figures 2b and 4e, the displacement of the peptide’s C-terminal anchor is a likely consequence of active displacement, allowing the collective strength of other specificity-determining interactions to be tested, resulting in a greater likelihood of dissociation for peptides with suboptimal anchors. The peptide-deficient state is often characterised by an internal hydrogen bond linking Asn77 to the effector sidechain, against which incoming peptides must compete. For HLA-B*44:05, our simulations suggest that being unable to hydrogen bond with either Asn77 or the penultimate carbonyl of the peptide, the sidechain of Trp147 flips outwards of the peptide binding groove, bringing the α helices closer together such that a hyper-closed conformation is adopted (figure 1d). However, this conformation is unlikely to be conducive for peptide binding, with breakage of the transient Tyr116-Asn77 hydrogen bond reorientating Trp147 such that a hydrogen bond can be shared with Asn77, regenerating a peptide receptive conformation. Our proposed mechanisms are supported by observations that peptides with suboptimal P9 were poorer competitors for binding to HLA-C*05:01 than to HLA-C*08:02. Interestingly, we observed that the two peptides that were most resistant to active displacement in HLA-C*05:01 (figure 5c) were the only peptides in cohort 1 with phenylalanine at position 1. Determining why peptides such as these can resist active displacement may allow the identification of superior vaccine immunogens. Our model is consistent with an experimentally validated systems model for quantitative antigen presentation in which the key step determining HLA-allotype dependency is the rate of transition between MHC-I structural intermediates (12, 51): a parameter that could relate to the rate (or probability) of active displacement among MHC-I allotypes.

Previous simulations of peptide-deficient MHC-I identified peptide binding groove plasticity (51), F-pocket conformational fluctuation (50) or F-pocket acidity (72) as determinants of independent peptide editing. Our simulations did not reveal differences in C or F pocket distances that correlated with the tapasin-dependence of the MHC-I allotypes in our panel (supplementary figure 3), indicating that global peptide binding groove dynamics do not underpin differences in tapasin-dependence. We believe that prior observations of correlations between tapasin-dependency and F pocket acidity or hydrophobicity are likely precursor observations of a very specific hydrogen bonding interaction that mediates active displacement and consequently independent peptide editing. Indeed, this may explain why mutations designed to increase the acidity of the F pocket are not always successful in increasing tapasin-mediated peptide editing of MHC-I allotypes (72).

Our analysis of the presentation of the KRAS-G12D neoantigen by HLA-C*08:02, but not by HLA-C*05:01, exemplifies how active displacement might influence the presentation of specific peptides in a physiologically relevant way. The position 77 and 80 sequence differences between HLA-C*05:01 and HLA-C*08:02 define two classes of HLA-C, known as C2 and C1 respectively, which form ligands for members of the killer-cell immunoglobulin-like receptor (KIR) family (55, 73). It is possible that the HLA-C active displacement mechanism, exemplified by HLA-C*05:01, may be generalisable to other HLA-C C2 allotypes, and explain why peptides with p9 alanine are more prevalent in HLA-C C1 subtypes (including HLA-C*08:02) immunopeptidomes than in HLA-C C2 subtypes (including HLA-C*05:01) (56, 57, 74). We further showed that a mutation designed to disrupt the independent editing mechanism had a modest impact on peptide presentation by HLA-B*44:05. This was evident at the level of individual peptides, and at the cell surface expression level. However, this disruptive mutation alone was not sufficient to abrogate independent peptide editing akin to the D116Y polymorphism defining the HLA-B*44:02/B*44:05 allotypes. Instead, this may reflect that the D116Y polymorphism changes peptide binding specificity (41, 42, 53) as well as the ability to disrupt the Asn77-peptide backbone hydrogen bond to mediate active displacement.

It is likely that the mechanisms by which tapasin and TAPBPR perform their peptide editing functions are extensions of this active displacement mechanism: structural and biophysical analyses indicate the binding of tapasin or TAPBPR deforms the MHC-I structure compared with that of a peptide-loaded MHC-I molecule (16, 17, 75–80). The alterations to the molecular structures are most apparent surrounding the peptide’s C-terminus, where the binding of tapasin and TAPBPR widen the α2-1 sub-helix and the opposing end of the α1 helix can be disordered (75) or deformed by the binding of calreticulin (16). The editing loop of TAPBPR, and by extension tapasin (16, 81) may also contribute to disrupt interactions between MHC-I and the peptide backbone in an analogous fashion to that described for independent peptide editing.

In summary, we identified a generalisable mechanism of tapasin-independent peptide editing by MHC-I allotypes using REST2 enhanced sampling molecular dynamics simulations which we have validated experimentally for single point mutants of HLA-B*44:05 and HLA-C*05:01. We view active displacement as a constitutive mechanism that facilitates selective dissociation of peptides by transient disruption of hydrogen bonds between MHC-I and the peptide backbone. However, the specific residues responsible for this hydrogen bond network may differ between allotypes, exemplified here by the differences between the displacement mediated by HLA-B*44:05 and HLA-C*05:01. This finding helps understand how minor sequence differences in the peptide binding groove can control peptide selection leading to dramatic differences in antigen presentation. If there is an equivalent internal, or intrinsic, hydrogen bond in the peptide empty state for every conserved hydrogen bond between peptide and MHC-I (such as the Tyr116-Asn77 or Asn77-Lys80 for HLA-B*44:05 or HLA-C*05:01 respectively), it follows that those MHC-I allotypes, or MHC-like molecules, with the greatest number of hydrogen bonding interactions between peptide and MHC-I might be expected to be the most stable in the peptide empty state. We hope these results encourage further investigation into localised active displacement mechanisms of specific MHC-I allotypes that drive antigen selection, determine immunopeptidome composition and subsequent T cell activation.

## Data availability statement

The datasets presented in this study are available upon request and will be made available in online repositories following peer review.

## Acknowledgements

The authors declare that financial support was received for the research and/or publication of this article. This work was funded by a Cancer Research UK program grant A28279 (awarded to TE). The authors thank Professor Mary Carrington and Dr Arman Bashirova for the kind provision of MHC-I expression levels measured in the absence of tapasin and published in ref (33). The authors acknowledge Max Quastel for critical reading of the manuscript.

## Ethical considerations

Ethical approval was not required for the studies on humans in accordance with the local legislation and institutional requirements because only commercially available established cell lines were used.

## Artificial intelligence statement

No potentially identifiable images or data are presented in this study. The author(s) declare that no Generative AI was used in the creation of this manuscript.

## Materials and methods

### Molecular Dynamics

Peptide was removed from peptide-loaded MHC-I crystal structures due to a lack of peptide-deficient MHC-I crystal structures. Simulations for HLA alleles were performed using Gromacs 2021.2 (82) patched with Plumed v2.7.3 (83) using the Amber14SB forcefield (84). Crystal structures were protonated using HTMD 2.1 (ProPKa3.1) (85–87). Peptides were deleted from the structure. Protonated alleles were solvated in a cubic box in TIP3P waters using a 1.2 nm edge distance and neutralised to 150 mM ionic concentration with Na^+^ and Cl^-^ ions using the Joung and Cheatham parameter set (88). Mutations were introduced using PyMol 2.5.2 mutagenesis tool. Energy minimisation was initially performed using steepest descent with 5000 steps and step size of 0.01 nm. Resultant structures underwent conjugate gradient minimization with 0.005 nm step size until reaching machine precision. NVT equilibration was performed for 100 ps using the Bussi-Donadio-Parrinello thermostat at 300 K at 0.1 ps time constant. NPT equilibration was subsequently performed for an additional 100 ps using the Berendsen barostat with 1 bar reference pressure at 2.0 ps time constant. Each simulation replica began from the NPT equilibrated final step. Production runs were performed in the NVT canonical ensemble using the Nosé–Hoover thermostat with 2.0 ps time constant and 300 K. Long-range electrostatics were calculated using particle mesh Ewald (PME) with a short-range real space summation cut-off at 1.0 nm. Lennard-Jones interactions were similarly cut off at 1.0 nm with dispersion corrections applied. Production simulations were run with REST2 enhanced sampling using 18 replicas spaced across 300-450 K. Bias potentials were applied to peptide binding groove facing residues 4-11, 23-36, 46-85, 93-101, 112-118, 122-126, 137-180 on the MHC-I heavy chain (89). All simulations were performed for a minimum of 150 ns per replica, with HLA-B*44 alleles extended to 300 ns.

### MHC-I allotype panel

HLA-B allotypes were selected that had at least a ten-fold difference in tapasin-dependence as measured in reference (33), differed by three amino acids or fewer, and for which there was a crystal structure of at least one allotype. These criteria identified five pairs: HLA-B*44:02/HLA-B*44:05; HLA-B*44:03/HLA-B*44:05; HLA-B*57:01/HLA-B*57:02; HLA-B*57:01/HLA-B*57:03 and HLA-B*58:01/HLA-B*58:02. We included only one pair involving HLA-B*44:05 (HLA-B*44:02/HLA-B*44:05) and HLA-B*57:01 (HLA-B*57:01/HLA-B*57:03) on account of their likely similarity to the HLA-B*44:03/HLA-B*44:05 and B*57:01/HLA-B*57:02 allotype pairs. This analysis was supplemented with three well studied HLA allotypes: HLA-B*08:01; HLA-B*27:05; and HLA-A*02:01. PDB structures used for each allele simulation are as follows: HLA-B*44:02 (3KPM), HLA-B*44:05 (3KPQ), HLA-B*57:01 (5VUE), HLA-B*57:03 (5VUE), HLA-B*58:01 (5VWJ), HLA-B*58:02 (5VWJ), HLA-B*08:01 (4PRN), HLA-B*27:05 (5IB1), HLA-A*02:01 (3PWL), HLA-C*05:01 (6JTO), and HLA-C*08:02 (6ULK).

### Computational Analysis

C-pocket distance was defined as the difference between centre of masses for Cα atoms of residues 67-73 and 150-156. F-pocket distance was defined as the difference between centre of masses for Cα atoms of residues 74-85 and 138-149. Hydrogen bonding interactions were calculated using an in-house python script based on the MDAnalysis distance_array function. Hydrogen bonding frequency was defined as percentage of frames with a polar group distance below a specified cut-off value (3.5 Å) to an Asn77 polar sidechain atom. Interactions excluded residues 76-78, 97, 123, 139-158, as these were presumed to be sterically incapable of interacting in the presence of bound peptide ligand or found to make productive contacts to Asn77 in peptide-bound crystal structures, as well as excluding backbone polar atoms.

### Synthetic Peptides

The sequences of the unlabelled competitor peptides are indicated in table 5. These peptides and the fluorescent tetramethylrhodamine (TAMRA) labelled peptides: EEFGK^TAMRA^AFSF and SAEPK^TAMRA^PLQL where K^TAMRA^ denotes TAMRA labelled lysine were synthesized by Syn Peptides (Shanghai, China). The following UV-labile conditional peptide ligands were utilized: SEIDTVAjY (HLA-B*44) and GADGVGjSL (HLA-C), where j represents 3-amino-3-(2-nitro)phenyl-propionic acid. The UV conditional peptides were synthesized by Peptide Synthetics (Fareham UK). All peptides were reconstituted in dimethyl sulfoxide (DMSO).

### MHC-I protein Production

Plasmids encoding HLA-B*44:05 and HLA-B*44:05-W147A have been described previously (42). The HLA-B*44:05-Y116F mutant was created by site directed mutagenesis using 5’-CGCGGGTATGACCAGTTCGCCTACGACGGCAAG-3’ and 5’-CTTGCCGTCGTAGGCGAACTGGTCATACCCGC-3’ primers. Plasmids encoding HLA-C0501, HLA-C0802, and the C0501 N77S and HLA-C0501 K80N mutants have been described previously (57). Recombinant peptide loaded MHC-I complexes were obtained by combining solubilised heavy chain inclusion bodies with solubilised human β_2_m inclusion bodies and either the SEIDTVAjY (HLA-B*44) or GADGVGjSL (HLA-C) UV labile peptide in 8 M urea, 50 mM MES pH 6.5, 0.1 mM EDTA. Refolding was initiated by 14-fold dilution with cold 100 mM Tris pH 8, 2 mM EDTA, 0.4 M l-arginine hydrochloride, 5 mM reduced glutathione and 0.5 mM oxidised glutathione added over three hours whilst stirring to achieve final concentrations of 1 µM heavy chain, 2 µM β2-microglobulin and 40 µM (HLA-B*44) or 25 µM (HLA-C) UV labile conditional peptide. Two days later, the protein mixture was concentrated and purified by size exclusion chromatography using a Superdex 200 packed 26/600 gel filtration column (Cytiva) and phosphate buffered saline.

### *In vitro* peptide competition experiments

Peptide competition experiments were prepared in PBS supplemented with 0.5 mg/ml bovine serum albumin (BSA) and a final concentration of 1.67% DMSO. 160 nM of the HLA-B*44 or 200 nM of the HLA-C MHC-I molecules were supplemented with 20x excess β2-microglobulin and exposed to 366 nm light for 20 minutes at 4 °C. Twenty µl of the UV exposed proteins were added to a 96 well microplate, with each well containing 40 µl of a titration (0 - 83.33 µM) of an unlabelled peptide competitor, 2 nM of either the EEFGK*AFSF (HLA-B*44) or SAEPK*PLQL (HLA-C) TAMRA labelled peptides. Samples were prepared in duplicate and incubated overnight at 25 °C. At least two independent experiments were performed. Fluorescence polarisation measurements were taken using an I3x (Molecular Devices) with rhodamine detection cartridge. Binding of TAMRA-labelled peptide is reported in milli polarisation units (mP) and is obtained from the equation: mP = 1000 x (S – G x P) / (S + G x P), where S and P are background subtracted fluorescence count rates (S = polarisation emission filter is parallel to the excitation filter; P = polarisation emission filter is perpendicular to the excitation filter and G (grating) is an instrument and assay dependent factor. IC50 values were calculated by performing non-linear regression in GraphPad Prism using the one phase decay model, with plateaus constrained to 50.

### *In vitro* indirect measurements of HLA-B*44 peptide-MHC-I complex half-lives

Indirect peptide dissociation experiments were conducted essentially as described in ref (90). Experiments were performed in PBS supplemented with 0.5 mg/ml BSA and a final concentration of 1.67% DMSO. 160 nM of HLA-B*44 molecules were supplemented with 20x excess β2-microglobulin and exposed to 366 nm light for 20 minutes at 4 °C, before being incubated with an equimolar concentration of each of the unlabelled peptides, or no peptide (no peptide control), overnight at 25 °C in a volume of 105.6 µl. The next day 48 µl was added to each well of a 96 well microplate, before 12 µl of 4 µM EEFGK*AFSF TAMRA labelled peptide was added to each well, and fluorescence polarisation measurements were periodically taken at 25 °C for ∼200 hours. Samples were prepared in duplicate. Each peptide was tested in at least two independent experiments. Peptide-MHC-I half-lives were calculated by performing non-linear regression in GraphPad Prism using the one phase association model, with plateaus constrained to the maximum polarisation that was measured in the no peptide control.

## Contributions

S.T wrote the manuscript and performed simulation work. T.E conceived the study. S.T, R.D and A.V.H generated experimental results. All authors read and contributed to the manuscript. A.V.H, J.W.E and T.E supervised the work and manuscript production. A.V.H and T.E contributed equally to this work and share senior authorship.

## Notes

### Competing Interest Statement

The authors have declared no competing interest.

## References

1. K. Dhatchinamoorthy, J. D. Colbert, K. L. Rock, Cancer Immune Evasion Through Loss of MHC Class I Antigen Presentation. Frontiers in immunology 12, 636568 (2021).

2. J. C. Crispin, G. C. Tsokos, Cancer immunosurveillance by CD8 T cells. F1000Research 9 (2020).

3. J. Robinson et al., IPD-IMGT/HLA Database. Nucleic acids research 48, D948–D955 (2020).

4. D. Gfeller et al., The Length Distribution and Multiple Specificity of Naturally Presented HLA-I Ligands. J Immunol 201, 3705–3716 (2018).

5. A. H. Bakker et al., Conditional MHC class I ligands and peptide exchange technology for the human MHC gene products HLA-A1, -A3, -A11, and -B7. Proc Natl Acad Sci U S A 105, 3825–3830 (2008).

6. P. J. Bjorkman et al., Structure of the human class I histocompatibility antigen, HLA-A2. Nature 329, 506–512 (1987).

7. A. T. Nguyen, C. Szeto, S. Gras, The pockets guide to HLA class I molecules. Biochem Soc Trans 49, 2319–2331 (2021).

8. B. Reynisson, B. Alvarez, S. Paul, B. Peters, M. Nielsen, NetMHCpan-4.1 and NetMHCIIpan-4.0: improved predictions of MHC antigen presentation by concurrent motif deconvolution and integration of MS MHC eluted ligand data. Nucleic acids research 48, W449–W454 (2020).

9. L. Zhang, K. Udaka, H. Mamitsuka, S. Zhu, Toward more accurate pan-specific MHC-peptide binding prediction: a review of current methods and tools. Briefings in bioinformatics 13, 350–364 (2012).

10. N. Bloodworth, N. R. Barbaro, R. Moretti, D. G. Harrison, J. Meiler, Rosetta FlexPepDock to predict peptide-MHC binding: An approach for non-canonical amino acids. PLoS One 17, e0275759 (2022).

11. M. Bonsack et al., Performance Evaluation of MHC Class-I Binding Prediction Tools Based on an Experimentally Validated MHC-Peptide Binding Data Set. Cancer Immunol Res 7, 719–736 (2019).

12. D. S. M. Boulanger et al., A Mechanistic Model for Predicting Cell Surface Presentation of Competing Peptides by MHC Class I Molecules. Frontiers in immunology 9, 1538 (2018).

13. M. J. Androlewicz, K. S. Anderson, P. Cresswell, Evidence that transporters associated with antigen processing translocate a major histocompatibility complex class I-binding peptide into the endoplasmic reticulum in an ATP-dependent manner. Proc Natl Acad Sci U S A 90, 9130–9134 (1993).

14. J. J. Neefjes, F. Momburg, G. J. Hammerling, Selective and ATP-dependent translocation of peptides by the MHC-encoded transporter. Science 261, 769–771 (1993).

15. J. C. Shepherd et al., TAP1-dependent peptide translocation in vitro is ATP dependent and peptide selective. Cell 74, 577–584 (1993).

16. A. Domnick et al., Molecular basis of MHC I quality control in the peptide loading complex. Nature communications 13, 4701 (2022).

17. A. Blees et al., Structure of the human MHC-I peptide-loading complex. Nature 551, 525–528 (2017).

18. B. Ortmann et al., A critical role for tapasin in the assembly and function of multimeric MHC class I-TAP complexes. Science 277, 1306–1309 (1997).

19. B. Sadasivan, P. J. Lehner, B. Ortmann, T. Spies, P. Cresswell, Roles for calreticulin and a novel glycoprotein, tapasin, in the interaction of MHC class I molecules with TAP. Immunity 5, 103–114 (1996).

20. P. A. Wearsch, P. Cresswell, Selective loading of high-affinity peptides onto major histocompatibility complex class I molecules by the tapasin-ERp57 heterodimer. Nat Immunol 8, 873–881 (2007).

21. M. Howarth, A. Williams, A. B. Tolstrup, T. Elliott, Tapasin enhances MHC class I peptide presentation according to peptide half-life. Proc Natl Acad Sci U S A 101, 11737–11742 (2004).

22. A. P. Williams, C. A. Peh, A. W. Purcell, J. McCluskey, T. Elliott, Optimization of the MHC class I peptide cargo is dependent on tapasin. Immunity 16, 509–520 (2002).

23. M. Chen, M. Bouvier, Analysis of interactions in a tapasin/class I complex provides a mechanism for peptide selection. EMBO J 26, 1681–1690 (2007).

24. L. H. Boyle et al., Tapasin-related protein TAPBPR is an additional component of the MHC class I presentation pathway. Proc Natl Acad Sci U S A 110, 3465–3470 (2013).

25. G. I. Morozov et al., Interaction of TAPBPR, a tapasin homolog, with MHC-I molecules promotes peptide editing. Proc Natl Acad Sci U S A 113, E1006–1015 (2016).

26. C. Hermann, J. Trowsdale, L. H. Boyle, TAPBPR: a new player in the MHC class I presentation pathway. Tissue Antigens 85, 155–166 (2015).

27. E. T. Spiliotis, H. Manley, M. Osorio, M. C. Zuniga, M. Edidin, Selective export of MHC class I molecules from the ER after their dissociation from TAP. Immunity 13, 841–851 (2000).

28. C. Howe et al., Calreticulin-dependent recycling in the early secretory pathway mediates optimal peptide loading of MHC class I molecules. EMBO J 28, 3730–3744 (2009).

29. A. Neerincx et al., TAPBPR bridges UDP-glucose:glycoprotein glucosyltransferase 1 onto MHC class I to provide quality control in the antigen presentation pathway. eLife 6 (2017).

30. L. Sagert et al., The ER folding sensor UGGT1 acts on TAPBPR-chaperoned peptide-free MHC I. eLife 12 (2023).

31. P. A. Wearsch, D. R. Peaper, P. Cresswell, Essential glycan-dependent interactions optimize MHC class I peptide loading. Proc Natl Acad Sci U S A 108, 4950–4955 (2011).

32. W. Zhang, P. A. Wearsch, Y. Zhu, R. M. Leonhardt, P. Cresswell, A role for UDP-glucose glycoprotein glucosyltransferase in expression and quality control of MHC class I molecules. Proc Natl Acad Sci U S A 108, 4956-4961 (2011).

33. A. A. Bashirova et al., HLA tapasin independence: broader peptide repertoire and HIV control. Proc Natl Acad Sci U S A 117, 28232–28238 (2020).

34. R. Greenwood, Y. Shimizu, G. S. Sekhon, R. DeMars, Novel allele-specific, post-translational reduction in HLA class I surface expression in a mutant human B cell line. J Immunol 153, 5525–5536 (1994).

35. S. M. Rizvi et al., Distinct assembly profiles of HLA-B molecules. J Immunol 192, 4967–4976 (2014).

36. J. Kaufman, Generalists and Specialists: A New View of How MHC Class I Molecules Fight Infectious Pathogens. Trends Immunol 39, 367–379 (2018).

37. Y. Shionoya et al., Loss of tapasin in human lung and colon cancer cells and escape from tumor-associated antigen-specific CTL recognition. Oncoimmunology 6, e1274476 (2017).

38. T. H. Hansen, M. Bouvier, MHC class I antigen presentation: learning from viral evasion strategies. Nat Rev Immunol 9, 503–513 (2009).

39. I. B. Harvey, X. Wang, D. H. Fremont, Molluscum contagiosum virus MC80 sabotages MHC-I antigen presentation by targeting tapasin for ER-associated degradation. PLoS Pathog 15, e1007711 (2019).

40. E. M. Bennett, J. R. Bennink, J. W. Yewdell, F. M. Brodsky, Cutting edge: adenovirus E19 has two mechanisms for affecting class I MHC expression. J Immunol 162, 5049–5052 (1999).

41. A. Kaur et al., Mass Spectrometric Profiling of HLA-B44 Peptidomes Provides Evidence for Tapasin-Mediated Tryptophan Editing. J Immunol 211, 1298–1307 (2023).

42. R. Darley et al., Evidence of focusing the MHC class I immunopeptidome by tapasin. Frontiers in immunology 16, 1563789 (2025).

43. P. Chappell et al., Expression levels of MHC class I molecules are inversely correlated with promiscuity of peptide binding. eLife 4, e05345 (2015).

44. S. Paul et al., HLA class I alleles are associated with peptide-binding repertoires of different size, affinity, and immunogenicity. J Immunol 191, 5831–5839 (2013).

45. A. Kosmrlj et al., Effects of thymic selection of the T-cell repertoire on HLA class I-associated control of HIV infection. Nature 465, 350–354 (2010).

46. F. Sieker, S. Springer, M. Zacharias, Comparative molecular dynamics analysis of tapasin-dependent and -independent MHC class I alleles. Protein Sci 16, 299–308 (2007).

47. F. Sieker, T. P. Straatsma, S. Springer, M. Zacharias, Differential tapasin dependence of MHC class I molecules correlates with conformational changes upon peptide dissociation: a molecular dynamics simulation study. Mol Immunol 45, 3714–3722 (2008).

48. M. A. Garstka et al., Tapasin dependence of major histocompatibility complex class I molecules correlates with their conformational flexibility. FASEB J 25, 3989–3998 (2011).

49. M. G. Mage et al., A structural and molecular dynamics approach to understanding the peptide-receptive transition state of MHC-I molecules. Mol Immunol 55, 123–125 (2013).

50. E. T. Abualrous et al., F pocket flexibility influences the tapasin dependence of two differentially disease-associated MHC Class I proteins. Eur J Immunol 45, 1248–1257 (2015).

51. A. Bailey et al., Selector function of MHC I molecules is determined by protein plasticity. Scientific reports 5, 14928 (2015).

52. O. Fisette, S. Wingbermuhle, R. Tampe, L. V. Schafer, Molecular mechanism of peptide editing in the tapasin-MHC I complex. Scientific reports 6, 19085 (2016).

53. D. Zernich et al., Natural HLA class I polymorphism controls the pathway of antigen presentation and susceptibility to viral evasion. J Exp Med 200, 13–24 (2004).

54. C. A. Peh et al., HLA-B27-restricted antigen presentation in the absence of tapasin reveals polymorphism in mechanisms of HLA class I peptide loading. Immunity 8, 531–542 (1998).

55. M. J. Sim et al., Canonical and Cross-reactive Binding of NK Cell Inhibitory Receptors to HLA-C Allotypes Is Dictated by Peptides Bound to HLA-C. Frontiers in immunology 8, 193 (2017).

56. S. Sarkizova et al., A large peptidome dataset improves HLA class I epitope prediction across most of the human population. Nature biotechnology 38, 199–209 (2020).

57. M. J. W. Sim et al., T cells discriminate between groups C1 and C2 HLA-C. eLife 11 (2022).

58. E. Tran et al., Immunogenicity of somatic mutations in human gastrointestinal cancers. Science 350, 1387–1390 (2015).

59. M. J. W. Sim et al., High-affinity oligoclonal TCRs define effective adoptive T cell therapy targeting mutant KRAS-G12D. Proc Natl Acad Sci U S A 117, 12826–12835 (2020).

60. E. Tran et al., T-Cell Transfer Therapy Targeting Mutant KRAS in Cancer. The New England journal of medicine 375, 2255–2262 (2016).

61. G. Kaur et al., Structural and regulatory diversity shape HLA-C protein expression levels. Nature communications 8, 15924 (2017).

62. M. J. W. Sim, et al., Innate receptors with high specificity for HLA class I-peptide complexes. Sci Immunol 8, eadh1781 (2023).

63. Z. Kosaloglu-Yalcin et al., The Cancer Epitope Database and Analysis Resource (CEDAR). Nucleic acids research 51, D845–D852 (2023).

64. H. Reimann et al., Identification and validation of expressed HLA-binding breast cancer neoepitopes for potential use in individualized cancer therapy. J Immunother Cancer 9 (2021).

65. D. K. Wells et al., Key Parameters of Tumor Epitope Immunogenicity Revealed Through a Consortium Approach Improve Neoantigen Prediction. Cell 183, 818–834 e813 (2020).

66. F. T. Ilca, L. Z. Drexhage, G. Brewin, S. Peacock, L. H. Boyle, Distinct Polymorphisms in HLA Class I Molecules Govern Their Susceptibility to Peptide Editing by TAPBPR. Cell reports 29, 1621–1632 e1623 (2019).

67. Y. Sun et al., Xeno interactions between MHC-I proteins and molecular chaperones enable ligand exchange on a broad repertoire of HLA allotypes. Sci Adv 9, eade7151 (2023).

68. A. Halenius et al., Human cytomegalovirus disrupts the major histocompatibility complex class I peptide-loading complex and inhibits tapasin gene transcription. J Virol 85, 3473–3485 (2011).

69. J. Chen et al., The downregulation of Tapasin in dendritic cell regulates CD8(+) T cell autophagy to hamper hepatitis B viral clearance in the induced pluripotent stem cell-derived hepatocyte organoid. J Med Virol 96, e29546 (2024).

70. T. Ogino et al., Association of tapasin and HLA class I antigen down-regulation in primary maxillary sinus squamous cell carcinoma lesions with reduced survival of patients. Clin Cancer Res 9, 4043–4051 (2003).

71. L. Sokol et al., Loss of tapasin correlates with diminished CD8(+) T-cell immunity and prognosis in colorectal cancer. J Transl Med 13, 279 (2015).

72. H. Lan et al., Exchange catalysis by tapasin exploits conserved and allele-specific features of MHC-I molecules. Nature communications 12, 4236 (2021).

73. R. Biassoni et al., Amino acid substitutions can influence the natural killer (NK)-mediated recognition of HLA-C molecules. Role of serine-77 and lysine-80 in the target cell protection from lysis mediated by "group 2" or "group 1" NK clones. J Exp Med 182, 605–609 (1995).

74. M. Di Marco et al., Unveiling the Peptide Motifs of HLA-C and HLA-G from Naturally Presented Peptides and Generation of Binding Prediction Matrices. J Immunol 199, 2639–2651 (2017).

75. Y. Sun et al., CryoEM structure of an MHC-I/TAPBPR peptide-bound intermediate reveals the mechanism of antigen proofreading. Proc Natl Acad Sci U S A 122, e2416992122 (2025).

76. C. Thomas, R. Tampe, Structure of the TAPBPR-MHC I complex defines the mechanism of peptide loading and editing. Science 358, 1060–1064 (2017).

77. J. Jiang et al., Crystal structure of a TAPBPR-MHC I complex reveals the mechanism of peptide editing in antigen presentation. Science 358, 1064–1068 (2017).

78. J. Jiang et al., Structural mechanism of tapasin-mediated MHC-I peptide loading in antigen presentation. Nature communications 13, 5470 (2022).

79. I. K. Muller et al., Structure of an MHC I-tapasin-ERp57 editing complex defines chaperone promiscuity. Nature communications 13, 5383 (2022).

80. A. van Hateren, T. Elliott, Visualising tapasin- and TAPBPR-assisted editing of major histocompatibility complex class-I immunopeptidomes. Curr Opin Immunol 83, 102340 (2023).

81. A. C. McShan et al., TAPBPR promotes antigen loading on MHC-I molecules using a peptide trap. Nature communications 12, 3174 (2021).

82. M. Abraham, Murtola, T, Schulz, R, Pall, S, Smith, JC, Hess, B, Lindhall E, GROMACS: High performance molecular simulations through multi-level parallelism from laptops to supercomputers. SoftwareX 1-2, 19–25 (2015).

83. M. Bonomi et al., Promoting transparency and reproducibility in enhanced molecular simulations. Nature Methods 16, 670–673 (2019).

84. J. A. Maier et al., ff14SB: Improving the Accuracy of Protein Side Chain and Backbone Parameters from ff99SB. Journal of Chemical Theory and Computation 11, 3696–3713 (2015).

85. S. Doerr, M. J. Harvey, F. Noé, G. De Fabritiis, HTMD: High-Throughput Molecular Dynamics for Molecular Discovery. Journal of Chemical Theory and Computation 12, 1845–1852 (2016).

86. C. R. Søndergaard, M. H. M. Olsson, M. Rostkowski, J. H. Jensen, Improved Treatment of Ligands and Coupling Effects in Empirical Calculation and Rationalization of pKa Values. Journal of Chemical Theory and Computation 7, 2284–2295 (2011).

87. M. H. M. Olsson, C. R. Søndergaard, M. Rostkowski, J. H. Jensen, PROPKA3: Consistent Treatment of Internal and Surface Residues in Empirical pKa Predictions. Journal of Chemical Theory and Computation 7, 525–537 (2011).

88. I. S. Joung, T. E. Cheatham, III, Determination of Alkali and Halide Monovalent Ion Parameters for Use in Explicitly Solvated Biomolecular Simulations. The Journal of Physical Chemistry B 112, 9020–9041 (2008).

89. S. Wingbermühle, L. V. Schäfer, Capturing the Flexibility of a Protein–Ligand Complex: Binding Free Energies from Different Enhanced Sampling Techniques. Journal of Chemical Theory and Computation 16, 4615–4630 (2020).

90. L. V. Brown, et al., De-risking clinical trial failure through mechanistic simulation. Immunother Adv 2, ltac017 (2022).

