## Supplementary figures 1, 2, 3, 4, 5 for "Atom-level mechanism of tapasin-independent peptide editing by Major Histocompatibility Complex class I molecules"

Supplementary Figure 1

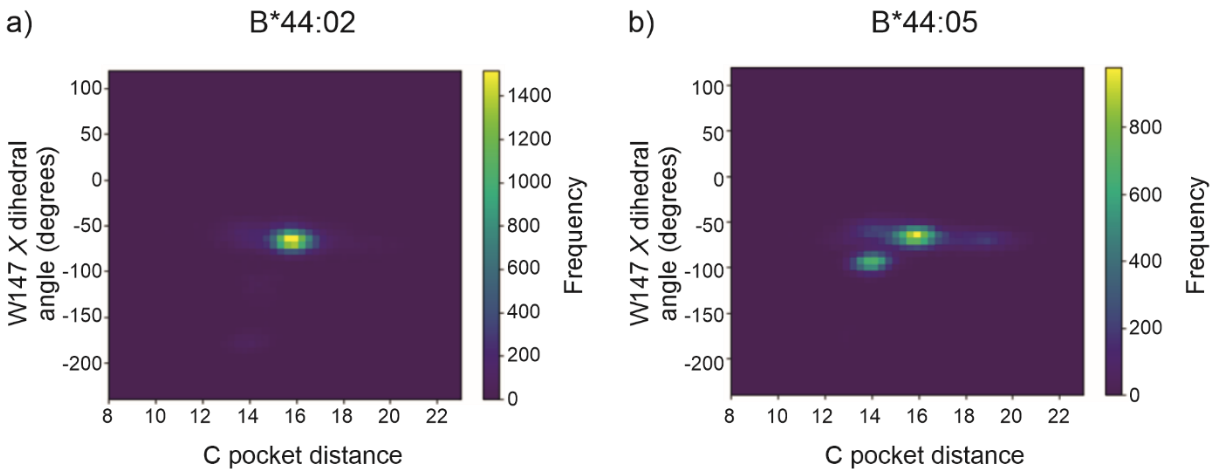


**Supplementary figure 1. Reorientation of Trp147 coincides with the hyper-closed conformation.**

a) Two-dimensional frequency histograms of the W147 sidechain rotational $\chi 1$ dihedral against the C-pocket distance for HLA-B*44:02.
b) Two-dimensional frequency histograms of the W147 sidechain rotational $\chi 1$ dihedral against the C-pocket distance for HLA-B*44:05.

Supplementary Figure 2

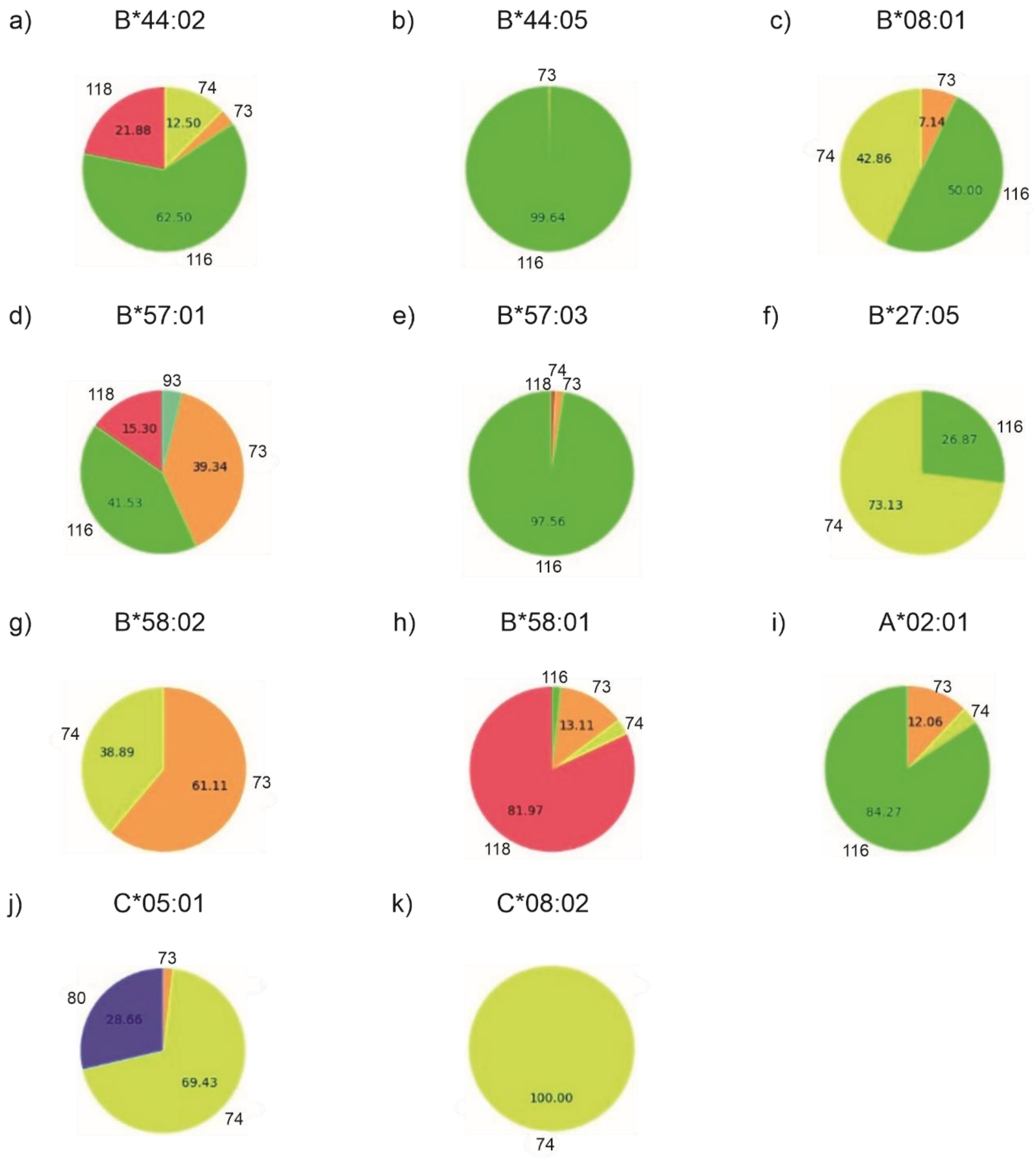


**Supplementary figure 2. Residue 77 hydrogen bonding interactions across REST2 simulations.**

Pie charts indicating the composition of residue 77 hydrogen bond interactions identified as in figure 2c. The HLA-B*44, HLA-B*57, HLA-B*58 and HLA-C allotype pairs are arranged in pairs with the independent self-editing allotype on the right.

Supplementary Figure 3

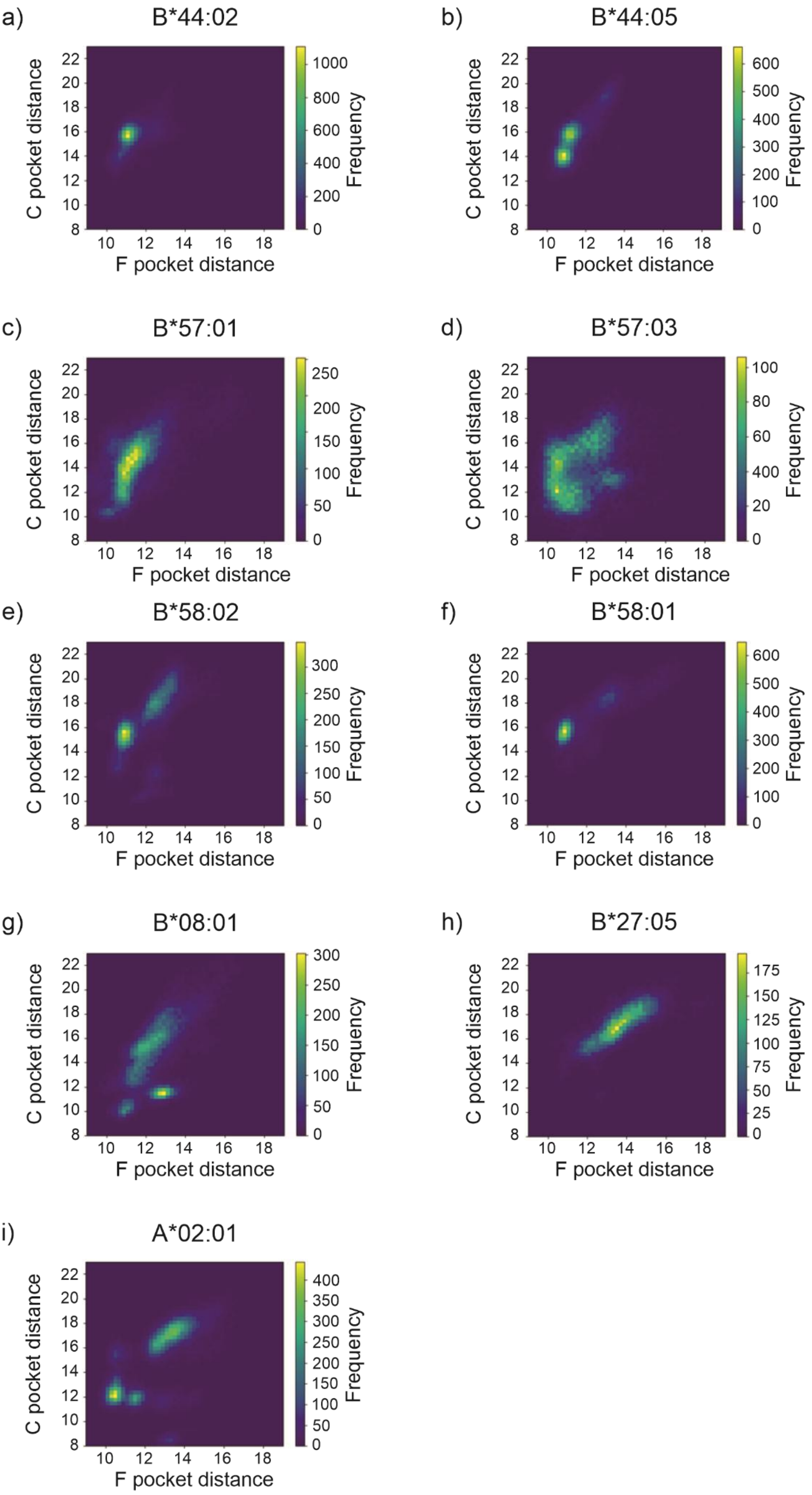


18

**Supplementary figure 3. Peptide binding groove dynamics does not correlate with tapasin-dependence.**

Two-dimensional frequency histograms of pocket distances from the indicated MHC-I allotypes, measured as in figure 1b. The HLA-B*44, HLA-B*57, HLA-B*58 and HLA-C allotypes (a-f) are arranged in pairs, with the independent self-editing allotype on the right.

Supplementary Figure 4

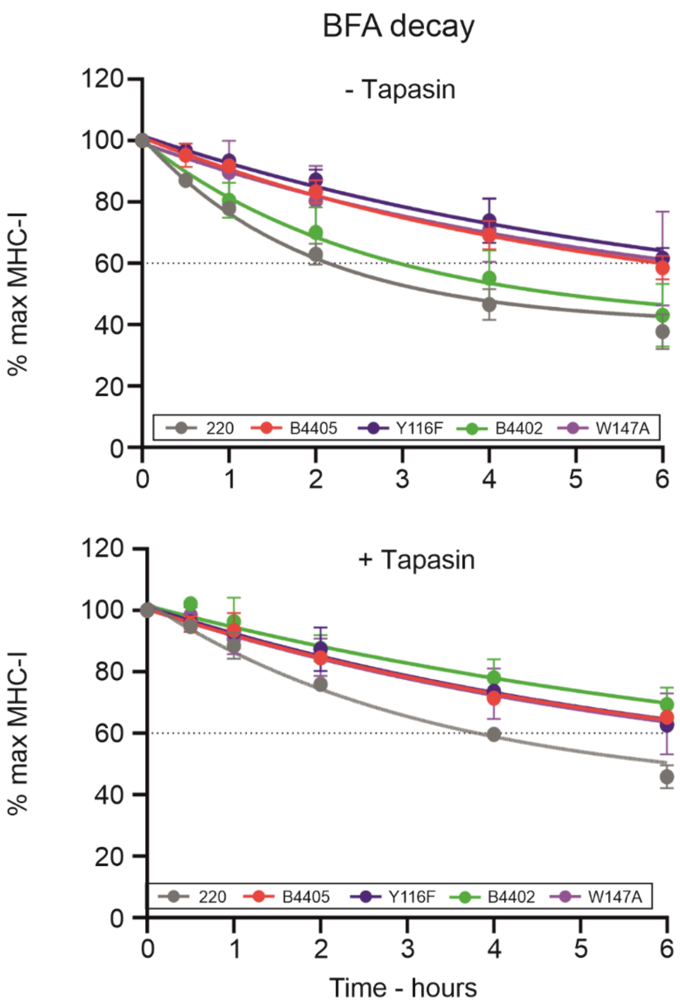


**Supplementary figure 4. HLA-B*44 cell surface stability measurements.**

Graphs depicting the cell surface expression levels of the indicated MHC-I molecules expressed in tapasin-deficient 721.220 cells (no tapasin, upper) or tapasin-reconstituted 721.220-tapasin cells (+ tapasin, lower) after culture in the presence of brefeldin A for the indicated length of time (x axes). MHC-I cell surface expression is plotted as a percentage of the initial cell surface expression level (y axes) and was measured by staining cells with W6/32 antibody and flow cytometry analysis. The graphs illustrate the mean fluorescent intensity measured in multiple experiments with vertical lines depict the standard deviation observed between three experiments.

Supplementary figure 5

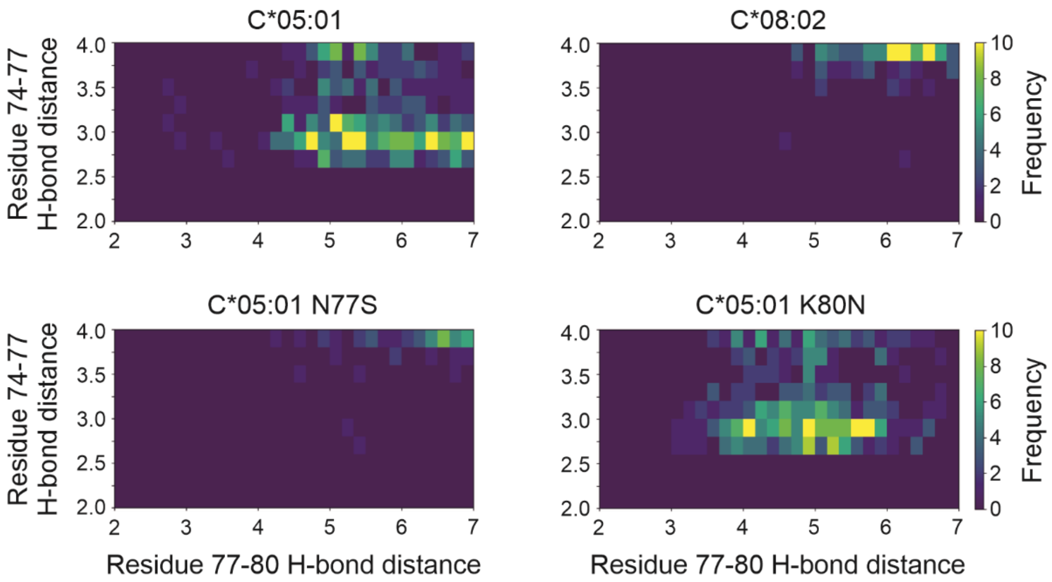


**Supplementary figure 5. Residue 77 hydrogen bonding in HLA-C.**

Identification of potential hydrogen bonds involving residue 77 in HLA-C from REST2 simulations. Panels show close-up views of the two-dimensional frequency histograms of the minimum distance between viable sidechain hydrogen bonding atoms plotted in figure 4b.
